# Activation of Wnt/β-catenin signalling by mutually exclusive *FBXW11* and *CTNNB1* hotspot mutations drives salivary gland basal cell adenoma

**DOI:** 10.1101/2024.10.21.619442

**Authors:** Kim Wong, Justin A. Bishop, Ilan Weinreb, Marialetizia Motta, Martin Del Castillo Velasco-Herrera, Emanuele Bellacchio, Ingrid Ferreira, Louise van der Weyden, Jacqueline M. Boccacino, Giovannina Rotundo, Andrea Ciolfi, Saamin Cheema, Rebeca Olvera-León, Victoria Offord, Alastair Droop, Ian Vermes, Michael Allgaeuer, Martin Hyrcza, Elizabeth Anderson, Katie Smith, Nicolas de Saint Aubain, Carolin Mogler, Albrecht Stenzinger, Mark J. Arends, Thomas Brenn, Marco Tartaglia, David J. Adams

## Abstract

Wnt signalling must be ‘just right’ to promote tumour growth. Basal cell adenoma (BCA) and basal cell adenocarcinoma (BCAC) of the salivary gland are rare tumours that can be difficult to distinguish from each other and other salivary gland tumour subtypes. Due to their rarity, the genetic profiles of BCA and BCAC have not been extensively explored. Using whole-exome and transcriptome sequencing of BCA and BCAC cohorts, we identify a novel recurrent *FBXW11* missense mutation (p.F517S) in BCA, that was mutually exclusive with the previously reported *CTNNB1* p.I35T gain-of-function (GoF) mutation. These driver events collectively accounted for 94% of BCAs. *In vitro*, mutant FBXW11 had a dominant negative affect, characterised by defective binding to β-catenin and the accumulation of β-catenin in cells. This was consistent with the nuclear expression of β-catenin observed in BCA cases harbouring the *FBXW11* p.F517S mutation and activation of the Wnt/β-catenin pathway. The genomic profiles of BCAC were distinct from BCA, with hotspot *DICER1* and *HRAS* mutations and putative driver mutations affecting PI3K/AKT and NF-κB signalling pathway genes. A single BCAC, which may represent a malignant transformation of BCA, harboured the recurrent *FBXW11* mutation. These findings have important implications for the diagnosis and treatment of BCA and BCAC, which, despite histopathologic overlap, may be unrelated entities.

## Introduction

Approximately 4-9% of all head and neck neoplasms are salivary gland tumours (SGTs), which include a wide range of benign and malignant entities that are further classified into distinct histopathological subgroups. Due to their rarity, overlapping presentation and the difficulty of their diagnosis, salivary gland basal cell adenoma and adenocarcinoma are of particular interest amongst these cancers. Cells of the intercalated duct have been proposed to be the primary cells of origin for BCA, a benign tumour^1^. BCAC, its presumed malignant counterpart, is thought to arise *de novo* in most instances, and in rare cases, arises from malignant transformation of BCA^2,3^. Of note, cases of BCA and BCAC have been linked to Brooke-Spiegler syndrome^4,5^, a rare autosomal dominant condition caused by germline disruption of the tumour suppressor gene *CYLD*.

The vast majority of salivary gland BCA and BCAC arise in the parotid gland^1,6–9^. BCA generally manifests as a slow-growing, well-defined, mobile, painless mass. In contrast, most BCAC are infiltrative and facial nerve involvement occurs in 25% of cases^10^. Local recurrence of BCAC has been reported in up to 50% of patients^11^ and occurs within 6-24 months after treatment^12^. Advanced BCAC is occasionally accompanied by cervical lymphadenopathy and distant metastases^13^. The clinical characteristics and natural history of these tumours highlight the need for a precise and prompt diagnosis for effective patient care, with effective surgical intervention being curative and metastatic spread of BCAC clinically challenging to manage.

Histopathologically, BCAs are well circumscribed or encapsulated showing tubulotrabecular, cribriform, membranous and/or solid growth patterns. Peripheral palisading of dark cells with luminal paler cells and ducts has been described in BCA, as have vesicular nuclei. The histological patterns of BCAC range widely and includes tubular, membranous, or solid growth patterns, with anaplasia being rare. As with BCA, nuclei are vesicular, while approximately a quarter of BCAC cases are accompanied by perineural and lymphovascular invasion. Diagnosis of BCA and BCAC may be aided by immunohistochemistry (IHC), through the assessment of β-catenin expression and markers of epithelial and myoepithelial differentiation; however, positivity for nuclear β-catenin has been found in both BCA and BCAC^14–16^. In other SGT subtypes, oncogenic, tumour-specific gene fusions have been identified^17–20^ and are increasingly used in combination with histology and IHC as a diagnostic aid^21,22^. Genetic studies of BCA have reported a recurrent missense activating mutation (p.I35T) in *CTNNB1*, the gene encoding the β-catenin protein^14,15,23^. Although the *CTNNB1* p.I35T mutation has not been found in BCAC, it has been reported in only a subset of BCA cases (37-80%)^24^, and, therefore, is of limited utility as a diagnostic tool. To date, genetic profiling of BCAC cases has revealed mutation of *PIK3CA*, *ATM*, *CYLD* and *NFKBIA*^14,23^, but given the rarity of this tumour type and the small number of genes profiled, the roles of these mutations in tumour development is unknown.

In this study, we sequenced the tumour and germline exomes and tumour transcriptomes of 32 cases of BCA and 11 cases of BCAC from multiple centres, an impressive collection given that these tumours represent approximately 1% of all salivary gland neoplasms and arise at a frequency of <1 per 100,000 individuals^25^. We identified and functionally characterised a recurrent somatic mutation in *FBXW11*, a novel driver event in BCA that is mutually exclusive with mutation of *CTNNB1*. In BCAC, we identified mutations at known hotspots in *HRAS*, *DICER1*, *PIK3CA* and somatic alterations that are predicted to promote tumourigenesis through diverse mechanisms. By examining somatic variants and copy number alterations, germline variants and gene fusions, we provide an unprecedented molecular portrait of these rare and diagnostically challenging head and neck tumours.

## Results

### Case selection and histopathological analysis

We collected formalin-fixed, paraffin-embedded (FFPE) samples from 75 patients, ascertained from six institutions across five countries. After sequencing and quality control, our cohort consisted of 65 cases; 55 cases with tumour and matched normal DNA and 10 tumour-only cases (Supplementary Table 1). Cases underwent a central pathology review by J.A.B. and I.W., who reviewed each case independently, arriving at a consensus diagnosis where their initial classification did not agree. This analysis, together with the genetic information from whole-exome and transcriptome sequencing, such as the presence of fusions diagnostic of other SGT types (described below), resulted in an analysis cohort of 43 cases (Table 1), each with matched normal tissue, including 32 BCAs, 9 BCACs and 2 salivary glands tumours with a differential diagnosis of BCAC and epithelial-myoepithelial carcinoma (BCAC/EMC) due to the presence of the oncogenic *HRAS* p.Q61R mutation (Supplementary Figure 1). Each case was from a different patient. IHC for β-catenin expression was not available for the majority of cases prior to pathology review by J.A.B and I.W. Data from transcriptome sequencing was available for all but 1 tumour. The BCAs were from 17 males (mean age of 65.7 years) and 15 females (mean age of 53.9 years), and the BCACs (including 2 BCAC/EMC) were from 6 males (mean age of 55.2 years) and 5 females (mean age of 55.4 years) (Table 1). The majority of the tumours were from the parotid gland. Representative hematoxylin and eosin (H&E) stains of a BCA and a BCAC from this study are shown in Supplementary Figure 2 and Supplementary Figure 3.

**Table 1:** Summary of salivary gland basal cell adenoma (BCA) and basal cell adenocarcinoma (BCAC) cases. Number of tumour DNA samples with a matched normal sample, number of RNA samples, age and sex of patients in the BCA and BCAC cohorts used in this study. The BCAC cohort included 2 BCAC with differential diagnosis of epithelial-myoepithelial carcinoma. T, tumour; N, normal; F, female; M, male. Tumours are primary tumours unless otherwise indicated.

| **Tumour type** | **DNA samples** | **RNA samples** | **Sex** | **Age, yrs (mean)** | **Site (Númer of tumours)** |
| --- | --- | --- | --- | --- | --- |
| BCAC | 11 T/N | 10 T | 5 F<br>6 M | 30-74 (55.2)<br>37-66 (55.4) F<br>30-74 (55.2) M | Parotid gland (7 primary, 1 recurrence)<br>Lung (2 metastases)<br>Sinonasal (1) |
| BCA | 32 T/N | 32 T | 15 F<br>17 M | 34-83 (60.1)<br>34-71 (53.9) F<br>36-84 (65.7) M | Parotid gland (29)<br>Submandibular gland (2)<br>Cheek (1) |

The 20 tumours (12 with matched normal) that were excluded from the final cohort of BCAs and BCACs included diagnostic mimics such as salivary gland pleomorphic adenomas (PAs), adenoid cystic carcinomas (ACCs), an epithelial-myoepithelial carcinoma (EMC), cases with differential diagnoses of myoepithelioma and carcinoma ex-PA (PA/ME/CXPA), and two cases that could not be definitively classified (NOS) (Supplementary Figure 1). Of note, several PAs were identified to have morphological features indicative of PA with a *HMGA2*::*WIF1* gene fusion^19^, which was subsequently confirmed following the analysis of the transcriptome sequencing data. Other cases had fusions involving *PLAG1*, which is commonly seen in PA/ME/CXPA^20^. Four of the 65 cases had *HRAS* hotspot mutations (three *HRAS* p.Q61R and one p.G13R; Supplementary Figure 1), which are frequently found in EMCs^26–28^ but have also been identified in other SGTs^23,29,30^. Due to this uncertainty, two BCAC cases with *HRAS* p.Q61R were retained for downstream analysis as BCAC/EMC, as described above, and a third, with *HRAS* p.G13R, was classified as a BCAC.

### Overview of somatic alterations in BCA and BCAC

Using whole-exome sequencing (WES) data, we identified somatic single nucleotide variants (SNVs), multinucleotide variants (MNVs) and insertion/deletion variants (indels) (Figure 1, Supplementary Figure 4 and Supplementary Table 2). The BCAC cohort included two lung metastases (PD56526a and PD56541a) and one recurrence (PD56535a; BCAC/EMC), which had higher mutation rates (1.5-2.4 mutations/Mb) than all but one primary BCAC/EMC (PD56546c; 7.5 mutations/Mb). (Figure 1, Figure 2 and Supplementary Table 3). The mutation rate of primary BCAC (0.22-7.5 mutations/Mb) was significantly higher than BCA (0.12-0.62 mutations/Mb) (Wilcoxon rank sum test, *p* = 0.0017), with median mutation rates of 0.56 and 0.32 mutation/Mb, respectively.

**Figure 1:**
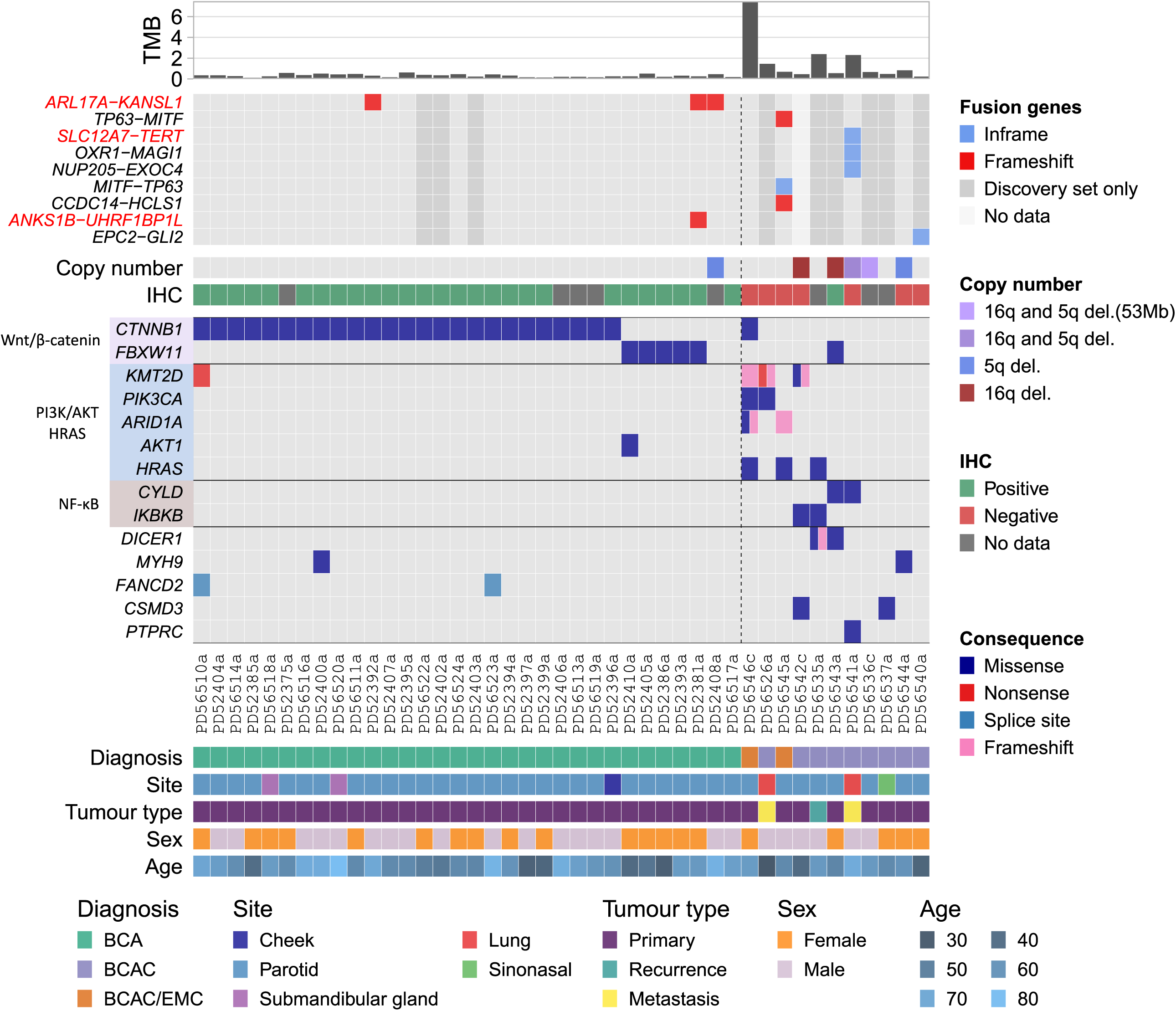
Overview of the salivary gland basal cell adenoma (BCA) and basal cell adenocarcinoma (BCAC) cohorts. BCA and BCAC cases are shown to the left and right of the dotted line, respectively. Genes shown in the oncoplot are genes known to be mutated in salivary gland tumours and/or are COSMIC Cancer Gene Census (CGC) genes and mutated in at least 1 BCA or BCAC. Genes that were mutated exclusively in the dMMR case, PD56546c, are not shown. TMB is the tumour mutation burden in mutations per megabase (Mb). Genes are grouped by pathway. TMB was calculated using mutations found in coding exons and 2 bp upstream and downstream of exon boundaries to account for splice site mutations. Fusion genes in red text are those found in the Trinity Cancer Transcriptome Analysis Toolkit human fusion library (see Methods). Samples marked as “Discovery set only” did not pass strict criteria for transcriptome sequencing quality control (see Methods) and may have a higher false discovery rate as a result. The copy number panel indicates the tumours that had copy number loss of chromosome arms 5q and/or 16q, which was significant in the BCAC cohort. BCAC/EMC is differential diagnosis of salivary gland basal cell adenocarcinoma and epithelial-myoepithelial carcinoma.

**Figure 2:**
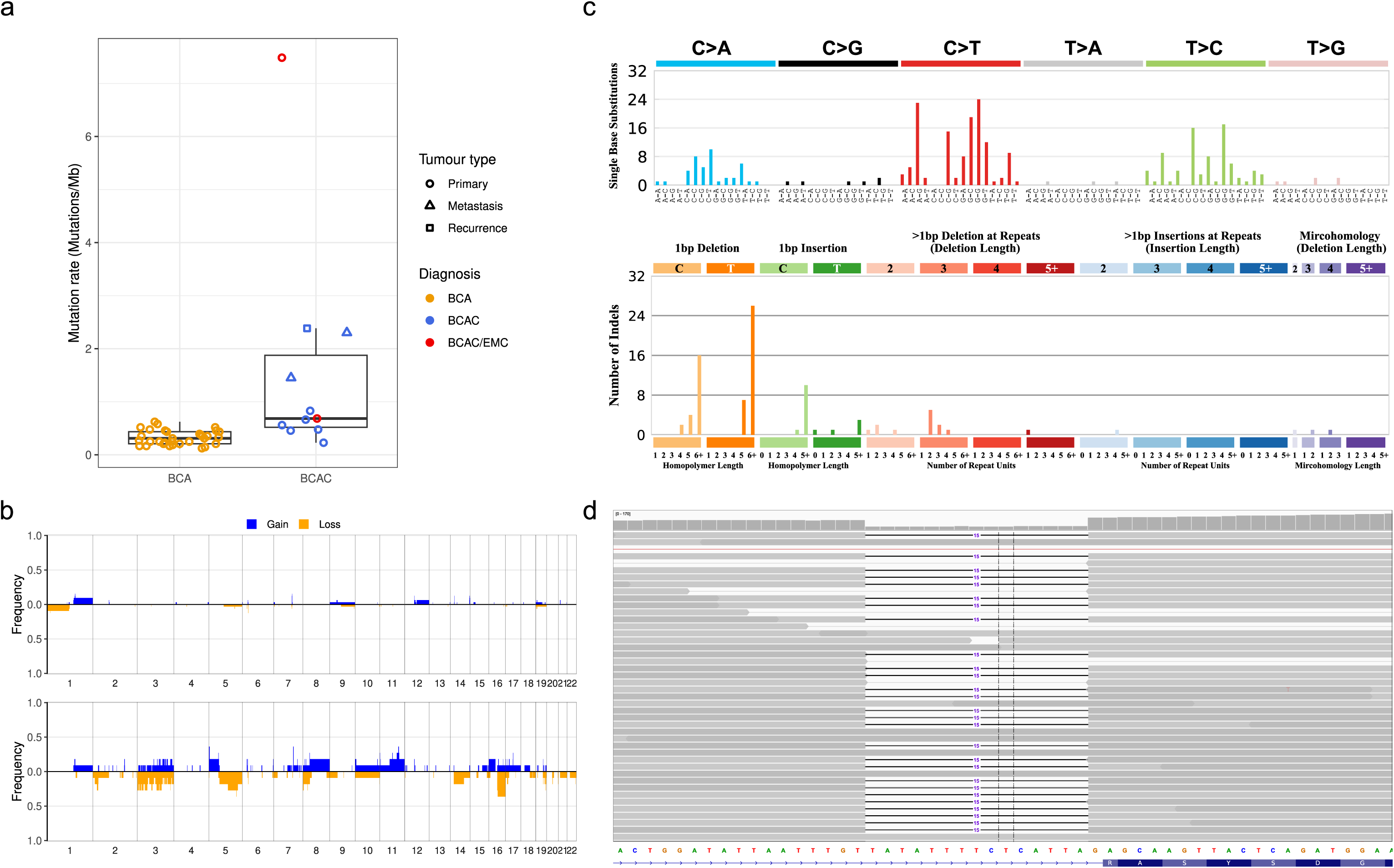
Tumour mutation burden, mutation spectra and recurrent copy number alterations. (a) Tumour mutation burden in salivary gland basal cell adenoma (BCA) and basal cell adenocarcinoma (BCAC). (b) Penetrance plot of somatic copy number gains and losses in BCA (top) and BCAC (lower). (c) The single base substitution spectrum (top) and indel spectrum (middle) from PD56546c. (d) PD56546c harboured a 15 bp somatic indel affecting the splice acceptor site in intron 4 or *MLH1* and loss of heterozygosity of chromosome 3p, leading to biallelic inactivation of *MLH1*.

The BCAC/EMC case (PD56546c) with an elevated mutation rate of 7.5 mutations/Mb (5.7 SNVs/Mb and 1.8 indels/Mb) was attributed to deficiency of mismatch repair (dMMR), as this case had SNV and indel mutational signatures found in tumours with dMMR and microsatellite instability (MSI) (COSMIC signatures SBS44, ID2 and ID7; Supplementary Figure 5) and biallelic inactivation of *MLH1* though somatic disruption of the splice acceptor site of intron 4 and copy number loss of chromosome 3q (Figure 2). No other COSMIC mutational signatures, other than ubiquitous signatures found in all tumour types (SBS1 and SBS5), were identified in either the BCA or BCAC cohorts.

Somatic copy number alterations (SCNAs) were infrequent in BCA, with a mean of 1.4% of the genome amplified or deleted (Figure 2). In contrast, a mean of 9.7% of the genome of BCACs were altered, with the highest proportion (29%) found in a lung metastasis (PD56541a). We identified a chromothripsis-like event (see Methods) on chromosome 3 of PD56545a, a BCAC/EMC, (Supplementary Figure 6), however, additional analyses, such as whole-genome sequencing, is required to confirm these events.

Unlike other SGTs, such as PA and ACC, recurrent gene fusions were not prevalent in BCA or BCAC (Figure 1 and Supplementary Table 4). High-confidence fusion genes (see Methods), identified through exome-based transcriptome sequencing from tumour samples, were identified in 3 BCAs and 3 BCACs. The only recurrent fusion identified was an inframe *ARL17A*::*KANSL1* fusion, which was present in 3 BCAs, that resulted from a ∼250kb deletion. Additional experiments are required to validate this fusion and determine its somatic status. One lung metastasis, PD56541a, harboured 3 gene fusions, including an inframe *SLC12A7*::*TERT* fusion. *SLC12A7*::*TERT* fusions have previously been reported in hepatocellular carcinoma^31^ and lung cancer^32^.

### A recurrent *FBXW11* p.F517S mutation is mutually exclusive with *CTNNB1* p.I35T in BCA

Previous studies involving a small number of BCAs have identified a recurrent p.I35T activating mutation in *CTNNB1*, which encodes the β-catenin protein^14,15,23^. In our cohort of 32 BCAs, *CTNNB1* was significantly mutated (25/32 or 78% of cases; *q* = 0.0; Benjamini-Hochberg method; see Methods), and remarkably all *CTNNB1* mutations were p.I35T (Figure 1 and Supplementary Table 5). This is in line with previous reports of *CTNNB1* p.I35T occurring in 37-80% of BCAs^24^. A *CTNNB1* p.D757N substitution in exon 15, which has not been reported in the COSMIC (v97) database, was found in the dMMR/MSI BCAC/EMC case PD56546c. However, it is unknown whether this mutation is functionally relevant; the majority of *CTNNB1* activating mutations are located in exon 3, which encodes serine/threonine phosphorylation sites that regulate β-catenin degradation^33^.

*FBXW11* was also identified as significantly mutated in BCA (5/32 cases; *q* = 4.0e-05; Benjamini-Hochberg method), and was mutually exclusive with mutation of *CTNNB1* (*q* = 0.012; discrete Benjamani-Hochberg method) (Figure 1). *FBXW11*, located on chromosome 5, encodes a 563 aa protein (Ensembl ID ENSP00000428753.2, encoded by the canonical transcript ENST00000517395.6) that is characterised by an *N*-terminal homodimerization domain, a central F-box region, and 7 tandemly arranged WD40 repeats at the *C*-terminus that constitute a WD repeat (WDR) domain (Figure 3). A recurrent missense mutation in exon 13 of *FBXW11*, p.F517S (NC_000005.10:g.171868777A>G; NM_001378974.1:c.1550T>C), was present in all 5 BCA with altered *FBXW11* (Figure 1), with one tumour harbouring an additional *in cis* mutation 28 bp downstream (NM_001378974.1:c.1578C>G; p.I526M) (Figure 3). The *FBWX11* p.F517S mutation was also found in a single high grade, malignant primary BCAC, PD56543a. The mutation was also present in the transcriptome sequencing reads from all cases (Figure 3). This mutation, which is predicted to be deleterious based on a SIFT score of 0 and its location within one of 7 WD40 domains (Figure 3), has not previously been reported in the ClinVar (release date 2023/01/21), dbSNP (v155) or COSMIC (v97) databases, though germline mutations affecting different residues located in the same domain have been reported to cause a syndromic neurodevelopmental disorder^34^. A literature search revealed a single report of this mutation in the OKAJIMA gastric cancer cell line^35^. Therefore, we used an orthogonal approach, PCR amplification, shotgun cloning, and Sanger sequencing, to confirm that the *FBXW11* mutation was indeed present in these cases and somatic (Figure 3c, Figure 3d and Supplementary Table 6).

**Figure 3:**
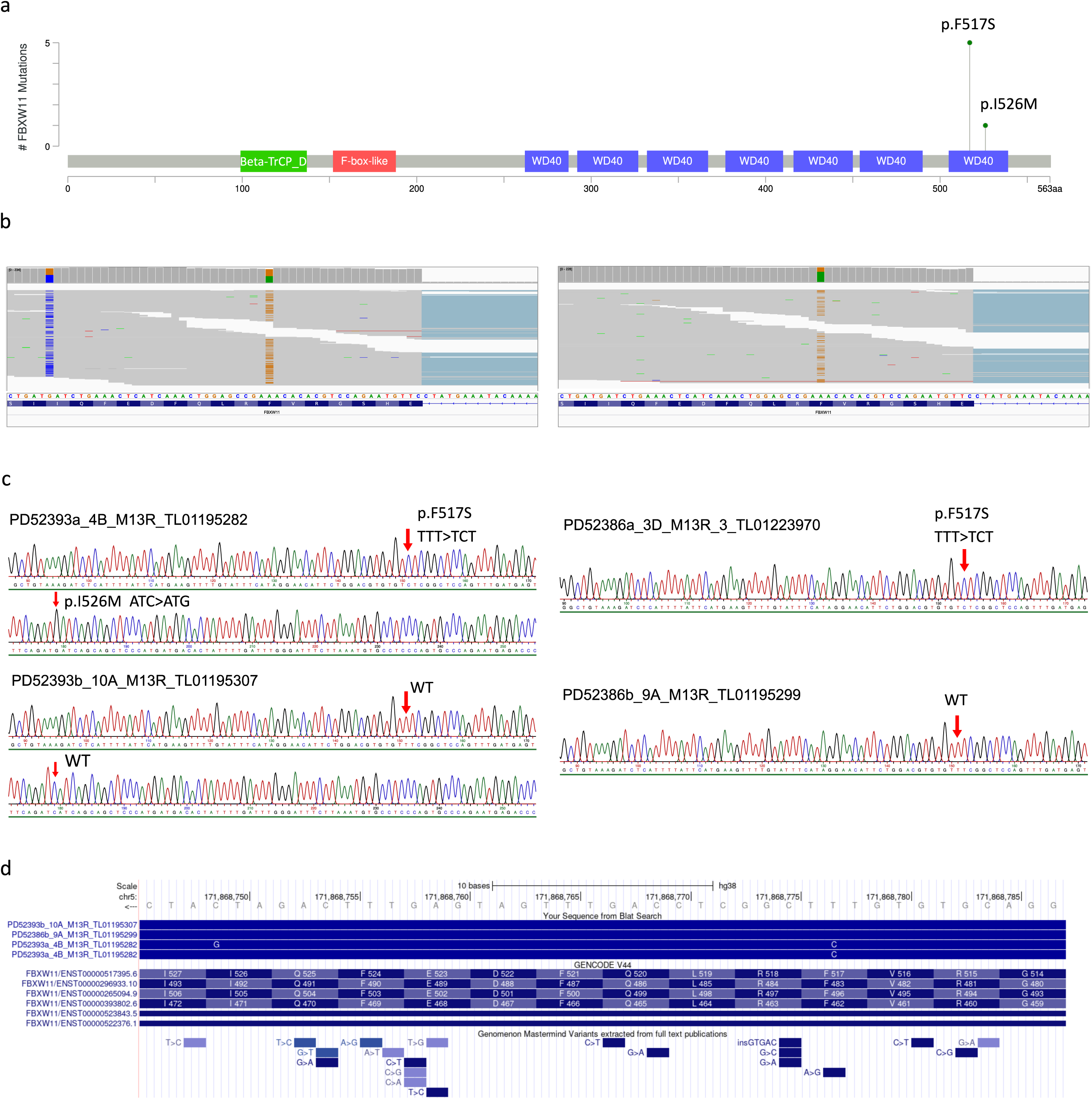
Validation of mutations in *FBXW11*. (a) The location and frequency of the p.F517S and p.I526M mutations in FBXW11. The amino acid position is shown on the x-axis, and coloured boxes represent protein domains. Beta-TrCP_D is the D domain of beta-TrCP; also shown are the location of an F-box domain and WD-40 repeats. (b) Examples of mutant alleles observed in tumour RNA-seq reads from PD52393a (left) and PD52386a (right). Tumour PD52393a had a p.F517S (NM_001378974.1:c.1550T>C) and a p.I526M (c.1578C>G) mutation. (c) Examples of traces generated from Sanger sequencing of clones derived from PCR amplification of *FBXW11* exon 13 in tumour (top) and matched normal (lower) DNA from the patients in (b). (d) Visualisation of the alignment of reads from (c), patient PD52393, in the UCSC Genome Browser, showing the location of mutant alleles relative to *FBXW11* exon 13. The A>G mutation at p.F517S (relative to transcript ENST00000517395.6) in the Genomenom Mastermind Variants track refers to the report of this mutation in the OKAJIMA cell line.

Although there was only a single occurrence of *FBXW11* p.F517S in the BCAC cohort, the difference in the proportion of BCA and BCAC cases with *FBXW11* p.F517S was not significant (*Χ*^2^ (1, *N* = 43) = 0.16, *p* = 0.70). This may indicate that the p.F517S mutation is not pathognomonic for BCA, however, the possibility that this case represents a malignant transformation of a BCA harbouring the *FBXW11* p.F517S mutation cannot be ruled out. The clinical history of the female patient indicated that she was evaluated at an emergency department (ED) after an 8-month history of having a lump on her cheek, which measured 1.7 cm by computed tomography, and again 6 years later, at which time the lump was 3.4 cm. The patient did not follow up for evaluation with an ear, nose and throat (ENT) specialist. A year later, the patient attended the ED with a 4 cm lump, where it was reported by the patient as having grown larger in recent years. The lump was removed after follow up with an ENT specialist. The given timeline is consistent with the hypothesis of BCAC arising from malignant transformation of BCA, however, because of the lack of an initial histopathological diagnosis, this cannot be confirmed. Molecular profiling of additional cases of BCAC is required to determine whether the *FBXW11* p.F517S mutation is pathognomonic of BCA or frequently mutated in both BCA and BCAC.

### *FBXW11* p.F517S reduces binding and polyubiquitination of β-catenin, leading to β-catenin accumulation

Activation of the Wnt/β-catenin pathway is a known mechanism of tumourigenesis that results from dysregulation of phosphorylation-mediated polyubiquitination and proteolysis of β-catenin, the protein encoded by *CTNNB1*. FBXW11 functions as a substrate adaptor of the SKP1-cullin-F-box (SCF) ubiquitin ligase complex, which catalyses phosphorylation-dependent ubiquitination of a wide array of substrates, including β-catenin^34,36–38^. Specifically, FBXW11 recognises and binds to phosphorylated β-catenin^36^. The SCF^FBXW11^ complex is a component of the β-catenin destruction complex involving AXIN, APC, CK1 and GSK3, which tightly regulates β-catenin in the absence of binding of Wnt ligands to membrane receptors^39^. In the presence of Wnt, β-catenin accumulates in the cytoplasm and is translocated to the nucleus where it activates Wnt target genes. Mutations in destruction complex components, such as those involving APC or AXIN, stabilise β-catenin, allowing its translocation to the nucleus^39^. The *FBXW11* p.F517S mutation is located in the last of seven WD40 repeats at the *C*-terminus of FBXW11 (Figure 3), which comprise a WDR substrate binding domain. Thus, we hypothesised that the *FBXW11* p.F517S mutation may affect binding to β-catenin.

To investigate the functional effects of the p.F517S mutation, we first performed molecular dynamics (MD) simulations on wild-type (WT) and F517S structures of FBXW11 (see Methods). F517 is in a solvent-exposed region in the WDR domain that is directly involved in β-catenin binding and *CTNNB1* D32 and G34 are key sites for the interaction of β-catenin and FBXW11^33,40^ (Figure 4a). F517 constrains a spatially close residue, R468, in a position that favours the formation of a salt bridge with D32 in β-catenin. The MD simulations predicted that the replacement of F517 with serine increases R468 mobility and the adoption of alternative conformations, causing this residue to move away from the β-catenin D32 residue (Figure 4a). Thus, this change was expected to disfavour the salt-bridge that stabilises the FBXW11-β-catenin complex and supports the hypothesis that the p.F517S substitution reduces the binding of FBXW11 to β-catenin.

**Figure 4:**
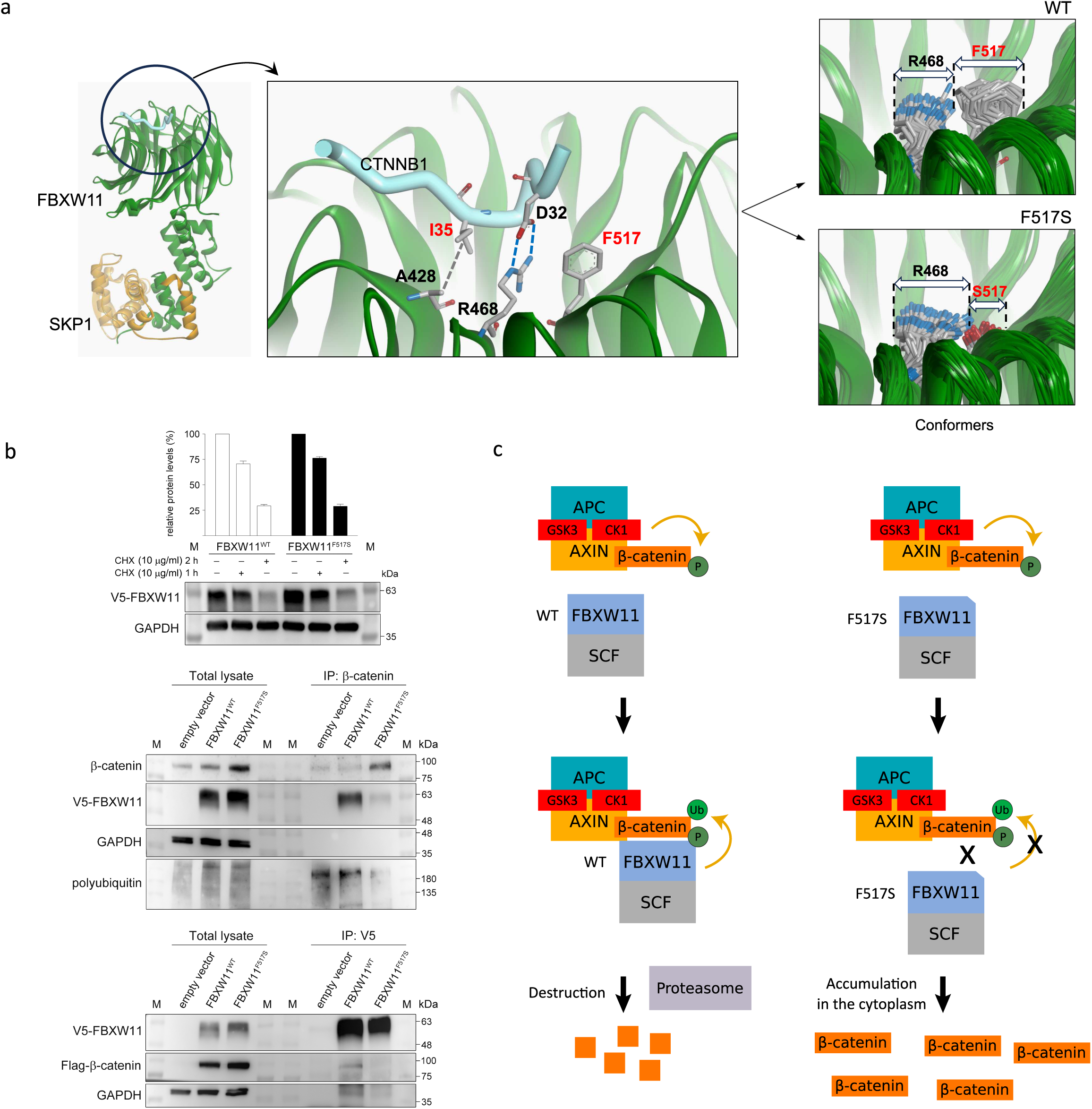
Functional consequences of the *FBXW11* p.F517S mutation. (a) The effects of the F517S substitution on the conformation of the FBXW11 region involved in β-catenin binding. Shown on the left is a complex of FBXW11-SKP1 with a β-catenin peptide phosphorylated on S33 and S37 (PDB ID 6WNX). The location of recurrently mutated sites at CTNNB1 I35 and FBXW11 F517 are indicated in red text. Substitution of phenylalanine with serine at position 517 increases the mobility of R468 and the adoption of alternative conformations (right, lower) compared to the wild type (WT) protein (right, top). (b) FBXW11^F517S^ is a stable protein (top panel). Western blot and relative quantitative analyses of V5-tagged FBXW11 protein levels in transfected COS-1 cells, basally and after cycloheximide (CHX) (10 μg/ml) treatment. GAPDH was used as loading control. The letter M denotes prestained protein markers. FBXW11^F517S^ has impaired ability to bind to β-catenin (middle and bottom panels). Lysates from transiently transfected HEK293T cells expressing V5-tagged FBXW11^WT^ or FBXW11^F517S^ proteins and serum-starved overnight were immunoprecipitated with anti-β-catenin antibody and assayed by western blotting using the antibodies indicated (middle panel). Note the reduced polyubiquitination of the endogenous immunoprecipitated β-catenin and the higher level of β-catenin in lysates from cells expressing FBXW11^F517S^. Lysates from transiently transfected HEK293T cells co-expressing FBXW11^WT^ or FBXW11^F517S^ proteins with Flag-tagged β-catenin and serum-starved overnight were immunoprecipitated with anti-V5 antibodies and assayed by western blotting using the antibodies indicated (bottom panel). Representative blots from one of three independent experiments are shown. (c) Left: FBXW11 is part of the Skp1-cullin 1-F-box (SCF) complex that regulates β-catenin levels by promoting its ubiquitination and subsequent proteasomal degradation. FBXW11 functions as substrate adaptor and is required for binding of phosphorylated β-catenin to the SCF complex, β-catenin phosphorylation is mediated by the destruction complex, which includes APC, AXIN, GSK3 and CK1. Right: the decreased binding of FBXW11^F517S^ to β-catenin inhibits the activity of the SCF complex.

Next, we performed *in vitro* experiments to confirm that the mutant FBXW11 causes a reduction in FBXW11-β-catenin binding and subsequent accumulation of β-catenin resulting from the reduction of β-catenin polyubiquitination. We first assessed the stability of the mutant protein in COS-1 cells transiently transfected to express either the FBXW11^WT^ or FBXW11^F517S^ protein (Figure 4b). This analysis demonstrated equivalent protein levels in cells treated with cycloheximide (CHX), indicating that the amino acid substitution did not impact proper protein folding or cause accelerated protein degradation (Figure 4b, top panel). Co-immunoprecipitation (co-IP) assays were performed using lysates collected from overnight-starved (serum-free Dulbecc’’s Modified Eagle Medium) HEK293T cells transiently expressing V5-tagged FBXW11^WT^ or FBXW11^F517S^ (Figure 4b, middle panel), or co-expressing either FBXW11^WT^ or FBXW11^F517S^ together with Flag-tagged β-catenin (Figure 4b, bottom panel). Binding of the FBXW11^F517S^ mutant to endogenous β-catenin was reduced compared with FBXW11^WT^, and, consistent with the model in which the inability of FBXW11^F517S^ to bind β-catenin impairs β-catenin degradation, we observed reduced polyubiquitination and consequent accumulation of endogenous β-catenin in cells expressing the mutant protein compared to FBXW11^WT^-expressing controls (Figure 4b, middle panel, and Figure 4c). Reduced binding of the FBXW11^F517S^ mutant to β-catenin was also confirmed using cells overexpressing Flag-tagged β-catenin (Figure 4b, bottom panel).

Given that *in vitro*, cells expressing FBXW11^517S^ accumulate β-catenin, we expected to observe nuclear expression of β-catenin in BCA with the *FBXW11* p.F517S mutation. Thus, we performed IHC staining for β-catenin on 21 BCA cases with *CTNNB1* p.I35T, 5 with *FBXW11* p.F517S and 1 case with neither mutation (Figure 1 and Figure 5). All cases we tested that harboured a mutation in either gene appeared similar, all having little or no membranous positivity with variable mild to moderate cytoplasmic staining and variable strong nuclear positivity staining, particularly in peripheral palisading dark cells around tumour cell islands. This pattern is consistent with increased accumulation of β-catenin in the BCA tumour cells. In contrast, normal salivary gland exocrine gland cells showed no nuclear staining. Similarly, all but one BCAC case for which IHC for β-catenin was performed (1/8) had membranous positivity and negative nuclear staining. The exception was the BCAC that harboured the *FBXW11* p.F517S mutation, which demonstrated patchy positive nuclear staining (Figure 5). A BCA with neither recurrent *CTNNB1* or *FBXW11* mutation was negative for nuclear staining. Other than *CTNNB1* and *FBXW11*, no other mutations in other Wnt/ꞵ-catenin signalling pathway genes (Supplementary Table 7) were detected in the BCA cohort. The discovery of the *FBXW11* p.F517S mutation and evidence that it activates the Wnt/β-catenin pathway therefore explains previous reports of β-catenin nuclear expression in BCA despite a lack of *CTNNB1* p.I35T mutation^14,15,23^.

**Figure 5:**
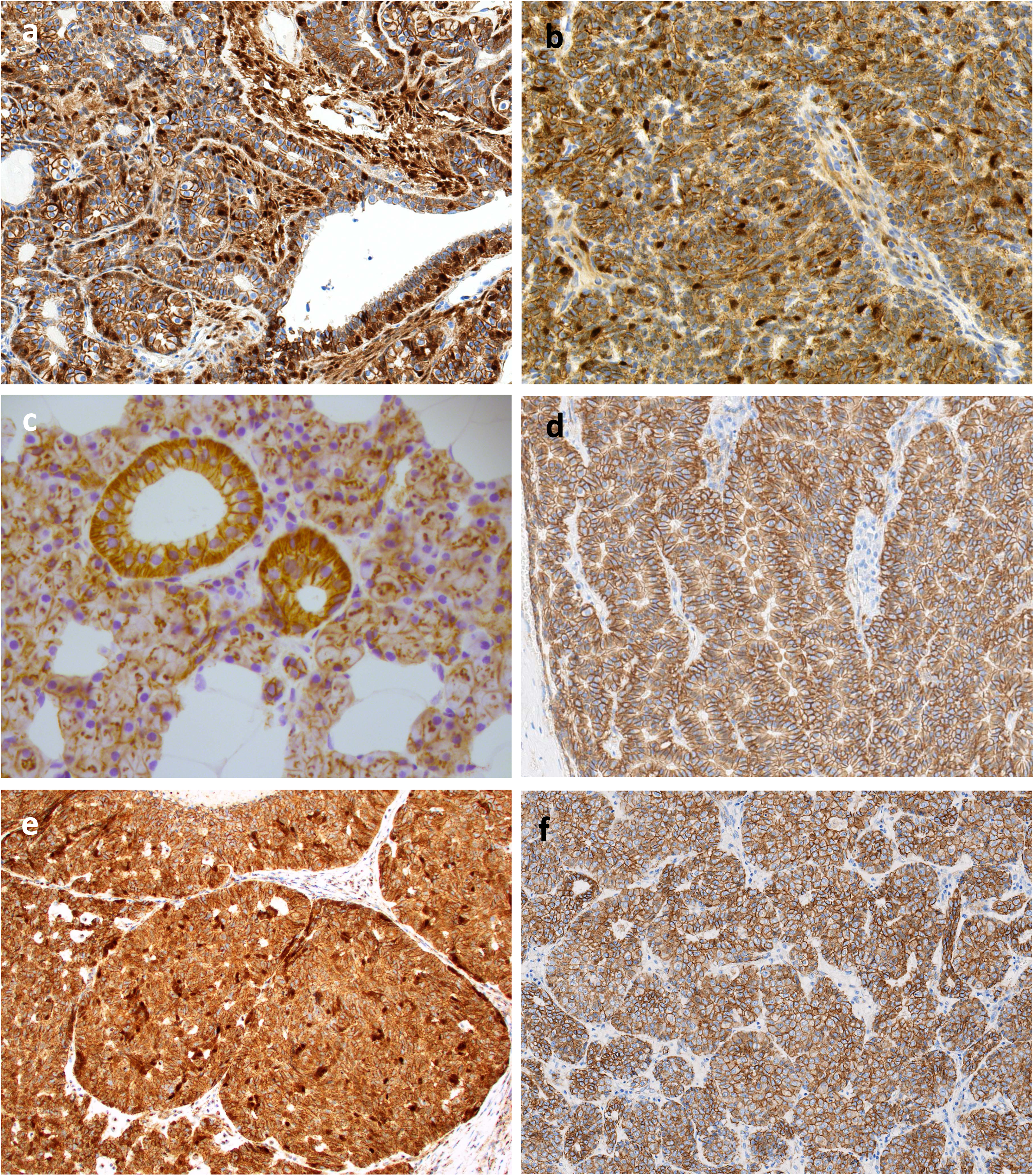
Immunohistochemical patterns of β-catenin in salivary gland basal cell adenoma (BCA) and basal cell adenocarcinoma (BCAC). (a) Case PD56510a, a BCA with a *CTNNB1* p.I35T mutation, demonstrating patchy nuclear β -catenin expression in both abluminal basal cells and stromal cells. (b) Case PD52381a, a BCA with a *FBXW11* p.517S mutation, exhibiting patchy nuclear β-catenin expression in a random pattern, along with focal nuclear expression in stromal cells. (c) Normal salivary gland exocrine gland cells from case PD52386a, with membranous β-catenin expression. (d) Case PD56517a, a BCA without a *CTNNB1* or *FBXW11* mutation, exhibiting membranous β-catenin expression. (e) Case PD56543a, a BCAC with a *FBXW11* p.517S mutation, showing patchy, random nuclear β - catenin expression. (f) Case PD56526a, a BCAC showing only membranous β -catenin expression.

The MD simulations also showed that the *CTNNB1* p.I35T substitution disrupts the hydrophobic interaction between amino acid residues I35 of *CTNNB1* and A428 of *FBXW11* (Figure 4a), and potentially perturbs the phosphorylation of adjacent residues S33 and S37. Phosphorylation at these sites, which are part of the β-catenin ubiquitination recognition site, is necessary for the targeting of β-catenin for proteasomal degradation^41,42^. This finding is consistent with β-catenin IHC nuclear positivity in samples harbouring the *CTNNB1* p.I35T mutation observed here and in other studies^14,15,23^.

Taken together, we can conclude that 30/32 BCA cases (93.7%) arose from activation of the Wnt/β-catenin signalling pathway, either through mutation of *CTNNB1* at p.I35T or *FBXW11* at p.F517S (Figure 1). Notably, this molecular mechanism does not represent a major driver event in BCAC.

### Copy number loss of chromosome 5q and 16q is frequent in BCAC

Amplification of oncogenes and deletion of tumour suppressors are known molecular mechanisms of tumourigenesis^43^. Copy number alterations have been profiled in common salivary tumours^44^, with distinct SCNAs found in different subtypes. To our knowledge, SCNAs have been described in one BCAC^45^ and one BCA^46^, respectively, using comparative genome hybridisation (CGH) and array CGH.

Using WES data, we generated allele-specific copy number (CN) profiles and identified significant recurrent CN alterations in the BCA and BCAC cohorts (Supplementary Table 8). In the BCA cohort, 1 focal deletion and 1 focal amplification were significantly recurrent (*q* = 0.022 and *q* = 0.0048, respectively; Benjamini-Hochberg method; Supplementary Table 8; see Methods); however, these regions harbour clusters of paralogous genes, making CN prediction from WES difficult.

Within our BCAC cohort, we identified 1 significantly recurrent focal deletion encompassing a 18 Mb region on chromosome 2 that included the tumour suppressor genes *DNMT3A* and *ASXL2* (Supplementary Table 8). Sequencing of additional BCAC cases may be required to provide statistical power. Chromosome arms 16q and 5q were significantly deleted (*q* = 8.6e-07 and *q* = 0.097, Benjamini-Hochberg method; see Methods) in 4/11 and 2/11 samples, respectively (Figure 1). Of note, 2 BCACs (one lung metastasis and one primary tumour) had biallelic inactivation of *CYLD*, which is located at 16q12.1, through loss of heterozygosity (LOH) and point mutation. Both point mutations were in the ubiquitin carboxyl-terminal hydrolase domain (Supplementary Figure 7), which is involved in deubiquitination of K63-linked TRAF proteins and negative regulation of NF-κB signalling^47^. *APC* and *FBXW11* are encoded on 5q, but no somatic or germline mutations in these genes were found and no other somatically mutated genes were shared amongst the samples with CN loss of 5q.

### Recurrently mutated genes and pathways in BCAC

While mutations in *PIK3CA* (p.H1047R), *NFKBIA* and deletion of *CYLD* have been identified in BCAC by either targeted panel sequencing or PCR and sequencing of individual genes^14,15,23^, we are the first to profile whole-exomes in BCAC. In our cohort, *KMT2D* and *HRAS* were identified as significantly mutated genes (*q* = 0.012 and *q* = 0.090, respectively; Benjamini-Hochberg method; Supplementary Table 5). *RPL22* was significant (*q* = 0.028; Benjamini-Hochberg method) when considering indel variants only. Our cohort of 11 BCACs included 2 cases of BCAC/EMC that each harboured *HRAS* p.Q61R hotspot mutations (Figure 1). The third case with an *HRAS* mutation (p.Q13R), PD56535a, was diagnosed as recurrent BCAC.

The histone transferase *KMT2D*, also known as *MLL2*, is a tumour suppressor and downstream target of the PI3K/AKT pathway^48^ that is frequently mutated in several tumour types^49^. In addition to the dMMR case PD56546c, two other cases harboured 2 *KMT2D* loss-of-function (LoF) mutations each; PD56526a, a lung metastasis with p.R3082Gfs*15 and p.C778*, and PD56542c, a primary tumour with p.C1408S (located within a PHD-finger and predicted to be deleterious with a SIFT score of 0) and p.F600Cfs*322 (Figure 1, Supplementary Figure 7 and Supplementary Table 2), which strongly suggests that biallelic inactivation of *KMT2D* may be a mechanism of tumourigenesis and/or progression in BCAC. Two cases also had hotspot GoF mutations in *PIK3CA*; the helical domain mutation p.E542K in PD56526a and the kinase domain mutation p.H1047R in PD56546c (Figure 1 and Supplementary Figure 7). A known recurrent *PIK3CA* GoF mutation, p.G118D, has been reported in one BCAC^14^. *PIK3CA* encodes the p110α subunit of the phosphoinositide 3-kinase (PI3K) pathway, which plays a role in cell survival and growth, and these hotspot mutations have been reported in a variety of tumour types^50^. Mutations were also observed in *ARID1A*, which can regulate PI3K/AKT pathway activity^51^.

In addition to the *FBXW11* p.F517S and *CTNNB1* p.D757N mutations in 2 cases described above, mutations in genes involved in regulation of Wnt/β-catenin and/or NF-κB signalling were found in 4/11 cases (36%) in the BCAC cohort, with mutations in *CYLD* (described above) and *IKBKB*. *CYLD* regulates the NF-κB and Wnt/β-catenin pathways, and loss of *CYLD* enhances Wnt/β-catenin signalling^47^. Biallelic inactivation of the NF-κB inhibitor alpha gene, *NFKBIA*, has been found in a BCAC^14^, however, we did not observe any non-silent somatic mutations affecting this gene. The inhibitor of nuclear factor kappa B kinase subunit beta gene, *IKBKB*, which encodes the IKKβ protein, was also mutated in 2 cases, with both harbouring missense mutations at codon 171 (p.K171T and p.K171M) within the protein kinase domain (Figure 1, Supplementary Figure 7 and Supplementary Table 2). Both variants are represented in the COSMIC database (COSV60598802 and COSV100645808, respectively) and have been previously identified in 3 lymphoid neoplasm samples each. Other substitutions at codon 171 have been shown to constitutively activate IKKβ, which in turn activates the NF-κB signalling pathway^52^. IKKβ has also been shown to phosphorylate β-catenin, which targets the protein for ubiquitination and degradation^53,54^.

*DICER1* is a tumour suppressor gene that encodes a ribonuclease involved in the processing of small RNAs, including those that play a role in RNA interference^55^. *DICER1* mutations were identified in 1 primary and 1 recurrent BCAC (Figure 1, Supplementary Figure 7 and Supplementary Table 2). The recurrent BCAC, PD56535a, harboured 2 somatic *DICER1* mutations, including the p.E1813D hotspot mutation located in the RNAse IIIb domain^55^ and a frameshift deletion at codon 1458 in the region encoding the RNAse IIIa domain. This biallelic hit of *DICER1* is similar to the two-hit pattern of *DICER1* mutations observed in tumours in patients with *DICER1* tumour predisposition syndrome, in which patients harbour two *DICER* mutations, a germline LoF variant and second mutation affecting the RNAse IIIb domain^56,57^. A second BCAC, PD56543a, had a p.S822T missense mutation located between the dsRNA-binding domain and PAZ domain. Neither patient had a pathogenic germline *DICER* variant.

### Germline and viral risk factors

Little is known about genetic risk factors for developing SGTs. Patients with Brooke-Spiegler syndrome (BSS), a rare, inherited condition attributed to germline disruptive variants in the *CYLD* gene develop tumours primarily on the face, scalp and neck, with cylindromas, spiradenomas and trichoepithelioma as the most common presentation, and rarely, SGTs^58,59^. Although somatic *CYLD* mutations were present in 2 BCAC, we did not identify any pathogenic germline variants in *CYLD* in either the BCA or BCAC cohort.

Interestingly, some salivary and mammary tumours share morphologic and genetic features^60^ and studies have suggested that a previous SGT is a risk factor for developing breast cancer^61–64^. Similarly, a link between germline *BRCA1* and *BRCA2* variants and an increased risk of developing SG tumours^65^ has been reported. In our BCAC cohort, *BRCA1* frameshift variants were present in the germlines of patients PD56541 (rs80357906) and PD56542 (rs80357971) (Supplementary Table 9 and Supplementary Figure 8). Both variants are reported in the ClinVar database (ClinVar Variant ID 17677 and 17667) as pathogenic breast and/or ovarian cancer susceptibility variants that have been reviewed by an expert panel. The tumours from these patients (one primary tumour and one lung metastasis) share somatic loss of 16q (Figure 1) but no other commonly mutated genes. Two BCA patients and 1 BCAC patient had germline *BRCA2* missense variants of uncertain clinical relevance. Pathogenic germline nonsense mutations in *PMS2* (rs63750451) and *MSH6* (rs267608094) were found in 2 patients (Supplementary Table 9 and Supplementary Figure 8), however, there was no indication of dMMR in the patients’ tumours, as the tumour mutation burden in these samples was < 1 mutation/Mb.

Human papillomavirus (HPV) has been identified in head and neck squamous cell carcinoma (HNSCC), with 70% of HNSCC of the oropharynx positive for HPV^66^. Consistent with previous findings that HPV is not associated with SGTs^66,67^, we did not identify HPV or other viruses associated with BCAC or BCA.

## Discussion

SGTs are rare, clinically heterogeneous and diagnostically challenging, with 39 tumour entities in the WHO 5th edition of salivary gland tumour classifications^3^. Here, we focused on BCAC, a rare malignant SGT, and BCA, which is considered its benign counterpart. To date, a limited number of genetic studies performed with small gene panels^14,23^ have examined these presentations. By using a multi-omics approach, we provide a comprehensive understanding of the different mechanisms driving BCAC and BCA. Our study also emphasises the challenge of diagnosing SGTs and the importance of molecular profiling, as a subset of cases originally diagnosed as BCA or BCAC were re-classified as other subtypes after consensus review of the histopathology in conjunction with genetic data.

The tumour suppressor *APC* is part of the destruction complex that regulates β-catenin, and in CRCs, *APC* inactivating mutations are almost always mutually exclusive with *CTNNB1* activating mutations^68^. Although we did not identify *APC* mutations, WES revealed a novel recurrent somatic driver mutation, *FBXW11* p.F517S (observed in 16% of BCAs), which was mutually exclusive with *CTNNB1* p.I35T (observed in 78% of BCAs). While *APC* mutations stabilise β-catenin by restricting the recruitment of kinases CK1 and GSK3 to the destruction complex^69^, we have shown that the p.F517S mutation directly reduces the binding of FBXW11 to β-catenin, resulting in defective ubiquitination and subsequent accumulation of β-catenin. This concurred with the observation that tumours with *FBXW11* p.F517S or *CTNNB1* P.I35T had positive β-catenin nuclear staining. Notably, in contrast to what has been observed in CRC, mutations in *CTNNB1* and *FBXW11* were limited to a single hotspot position in each gene. While sequencing of additional BCACs is required to confirm the frequency of *FBXW11* p.F517S, if this mutation is exclusive to BCA (and rare cases of BCAC that have arisen from BCA), our finding suggest that genetic testing for *CTNNB1* p.I35T and *FBXW11* p.F517S may be used to confirm nearly 94% of BCA cases helping to distinguish BCA from other mimickers.

In line with previous findings, the genetic profiles of BCACs were more varied and complex than BCAs. The lack of common driver genes and mutations, other than a single BCAC with *FBXW11* p.F517S, confirms that most BCACs arise *de novo* rather than from malignant transformation of BCA. In this study, *CYLD* mutations were found in only 2 of 11 BCAs and none of the 32 BCACs. A previous study suggested that 36% of BCAs and 29% of BCACs had nonsense mutations in *CYLD*^70^, however, because only 8 of the 45 BCAs and BCACs used in the previous study were paired with matched normal tissue^70^, the variants found may potentially be germline variants. In addition to *CYLD*, we identified mutations in *IKBKB*; both genes play roles in the Wnt/β-catenin and NF-κB signalling pathways. However, nuclear expression of β-catenin was negative in cases with these mutations, which suggests that the NF-κB pathway, rather than the Wnt/β-catenin pathway, might be activated via functional dysregulation of *CYLD* and *IKBKB* in BCAC. Previous studies have observed nuclear β-catenin expression in subsets of both BCAC and BCA with a wide range of nuclear positivity rates^14,16,71^. However, nearly all BCAs and only 1 of 8 BCAC cases we tested was positive, all of which could be attributed to the presence of either the *CTNNB1* p.I35T or *FBXW11* p.F517S mutation. The inconsistent findings amongst studies strengthens the argument that a combination of tools including histopathology, IHC and genetic profiling can aid diagnosis of SGTs, particularly rare subtypes.

This study is the first to profile somatic CN alterations in BCA and BCAC using NGS. Five of 11 BCACs had loss of one or both of chromosome arms 5q and 16q. This is also frequent in a subset of breast cancers^72^ and numerous studies have attempted to elucidate whether specific tumour suppressor genes are the target of 16q loss^73^. In BCAC, we found somatic biallelic inactivation of *CYLD* was a consequence of 16q loss in 2 of 4 cases, implicating *CYLD* as the target of 16q loss in BCAC.

Our analysis of BCAC has revealed several areas for further investigation. It would be of interest to determine the frequency of dMMR in BCAC, as these patients may benefit from immunotherapy in the metastatic setting. We also identified *KMT2D* as a novel candidate driver of BCAC and several candidate driver genes in the PIK3K/AKT and NF-κB pathways. Further exploration into the association between breast cancer and BCAC may be of relevance, given that 2 of 11 BCAC patients in this study had LoF *BRCA1* germline variants and previous studies have suggested a higher risk for breast cancer in patients with SGTs. Deconvolution of the roles of these pathways and genes will benefit patients, as treatment can be informed by genetic profiling of tumours.

This is the first study using whole-exome and transcriptome sequencing to explore the mutational landscape of BCA and BCAC. We have identified and functionally characterised a novel, recurrent driver mutation *FBXW11* p.F517S, which is mutually exclusive with the Wnt/β-catenin activating mutation *CTNNB1* p.I35T in BCA. The mutant FBXW11 displayed defective β-catenin binding, leading to the stabilisation of β-catenin, thus, serving as an alternative mechanism of Wnt/β-catenin signalling activation. We have also shown that BCAC involves several driver genes and pathways, indicating that most BCACs arise *de novo* and that BCA and BCAC may be unrelated entities. Importantly, our work has highlighted that molecular profiling, along with histopathology and IHC, can be used for more accurate diagnosis of SGTs.

## Methods

### Sample acquisition and nucleic acid isolation

Cases consisted of formalin-fixed, paraffin-embedded (FFPE) tissues that had been collected as part of routine diagnostic procedures with the patient’s consent. Ethical approval for the use of these samples and associated data was obtained by a local committee at the institution of origin and via Research Governance at the Wellcome Sanger Institute. Representative haematoxylin & eosin (HE)-stained sections of all FFPE tissue blocks were independently reviewed by two consultant pathologists (I.F. and T.B.) to confirm diagnoses and to identify areas for sampling (tumour and normal, where possible). All tumour and normal tissue samples were obtained as either 1 mm diameter cores or as unstained 10-micron thick tissue sections attached to glass slides (from which the tumour and normal areas were manually macro-dissected). Genomic DNA and RNA was extracted from the tumour samples (with genomic DNA only extracted from normal samples) using the AllPrep DNA/RNA FFPE Kit (Qiagen), according to the manufacturer’s instructions.

### DNA sequencing, read alignment and quality control

Sequencing libraries were prepared from FFPE-extracted DNA using a ‘NEB Ultra II RNA custom kit’ on an Agilent Bravo WS automation system. Unique dual index tags were appended, and the samples were amplified by PCR using the KAPA HiFi Kit (KAPA Biosystems) for a minimum of eight cycles. The libraries were quantified using the Accuclear dsDNA Quantitation Kit (Biotium), pooled (8-plex) in an equimolar fashion and hybridized overnight with SureSelect Human All Exon V5 baits (Agilent). The multiplexed samples were paired-end sequenced using the NovaSeq 6000 platform (Illumina) to generate 101 bp reads.

Sequencing reads were aligned to the GRCh38 reference genome, using `BWA-MEM`^74^ and PCR duplicates from the Binary Alignment Map (BAM) file were marked using the `samtools` (v1.14)^75^ `markdup` function with parameters `–mode s –S –include-fails`. Matched tumour-normal sample concordance, as well as cross-individual contamination, was assessed using `Conpair` (v0.2)^76^. After excluding samples with quality issues such as having less than 80% of the bait capture regions with a minimum of 20X coverage, matched tumour-normal genotype concordance <60%, or cross-individual contamination >5% and selecting one sample per tumour (where applicable), there were 48 cases diagnosed as BCA (including 7 cases without matched normal DNA available) and 17 samples diagnosed as BCAC (including 3 cases without matched normal DNA available). Given that histopathological diagnosis of salivary gland tumours can be challenging, the diagnoses for samples passing quality control were then independently re-reviewed by two specialist head and neck pathologists (J.A.B and I.W.). Slide images were reviewed, along with genetic data, which included fusion genes information associated with diagnostic mimics (for example the *NFIB*::*MYB* fusion in ACC and the *HMGA2*::*WIF1* fusion in PA) and somatic variants in genes previously described in BCA or BCAC, including *CTNNB1*, *PIK3CA* and *HRAS*. In cases where the diagnosis differed after independent review, a consensus diagnosis was derived after a joint review. This detailed case review resulted in a cohort of 11 BCAC and 32 BCA, each with matched normal DNA.

### Exome-capture transcriptome sequencing, read alignment and quality control

FFPE-extracted RNA samples were reverse-transcribed and sequencing libraries prepared using the NEBNext Ultra II Directional RNA Library Prep kit (New England Biolabs) according to the manufacturer’s instructions. Unique dual index tags were appended, and the samples were amplified by PCR using the KAPA HiFi HotStart ReadyMix PCR Kit (Roche) for a minimum of 16 cycles. Libraries were quantified using the Accuclear dsDNA Quantitation Kit (Biotium), pooled (8-plex) in an equimolar fashion and hybridized overnight with the SureSelect Human All Exon V5 baits (Agilent). The multiplexed samples were paired-end sequenced using the NovaSeq 6000 platform (Illumina) to generate 101 bp reads. Reads were aligned using `STAR` (v2.5.0c15)^77^ against the GRCh38 human reference genome using Ensembl release v103 gene annotations. Expression levels were assessed by counting reads using `HTseq` (v0.7.2)^78^ with the appropriate stranded parameter and subsequently transformed into transcripts per million (TPM) values. Data quality was assessed by running RNA-SeqQC 2^79^ and assessing the total number of counts obtained per sample. The criteria to select samples for a high-quality analysis cohort are provided in the Supplementary Methods. A total of 29 BCA and 4 BCAC RNA sequencing datasets passed these criteria. For consensus review of cases (described above), all cases for which sequencing data was available were used for fusion discovery (32/32 BCA and 10/11 BCAC). We refer to this as the “discovery set” of samples.

### Identification and annotation of somatic variants

Somatic point mutations were identified using `cgpCaVEMan` (v1.15.2)^80^. The parameters used and input files are described in the Supplementary Methods. Variant flagging and annotation were not performed initially. Instead, adjacent, *in cis* SNVs called with `cgpCaVEMan` were evaluated using `SmartPhase` (v1.2.1)^81^ and `casmsmartphase` (v0.1.8; https://github.com/cancerit/CASM-Smart-Phase) to identify MNVs. Variants were then flagged using the `cgpcavemanpostprocessing` (v1.10)^80^ `cgpFlagCaVEMan.pl` utility using the ‘WXS’ mode for exomes. The parameters and flagging rules used are described in the Supplementary Materials.

Indels on the autosomes and chromosomes X and Y were identified using `cgpPindel` (v.3.10.0)^82^. A simple repeats file for GRCh38, generated using the UCSC Table Browser, and a list of regions to exclude due excessive high depth of coverage, were used as inputs. Soft flag FF017 was used, but variants were not hard filtered based on this flag. Additional details, including a description of parameters, flags and input files, are available in the Supplementary Methods.

The Ensembl release v103 Variant Effect Predictor (`VEP`)^83^ was used to predict the consequences of SNVs, MNVs and indels on proteins. The canonical transcript, as defined by Ensembl, was used to determine the variant consequence. `VEP` was also used to add custom annotations from the COSMIC (v97)^84^, ClinVar (update 20230121)^85^, gnomAD (v3.1.2)^86^ and dbSNP (v155)^87^ databases.

Common SNPs, defined as variants with a minor allele frequency of at least 0.01 in the total population in the gnomAD database (v3.1.2) or the 1000 Genomes Phase 3 dataset (as indicated by dbSNP v155), were excluded as germline variants. `cgpPindel` calls were further refined by retaining variants with VAF >= 0.1, multinucleotide variants of length 3 bp or less, and variants with both the reference allele and alternative allele are <= 25 bp in length or more but both alleles are not >10 bp. Finally, variants within 100 bp of exome targeted regions and not flagged by `cgpCaVEMan` or `cgpPindel` filters were taken forward for further analysis.

Manual inspection of read alignments at recurrently mutated sites in *CTNNB1* and *FBXW11* was performed to minimise false negative calls at these sites due to the presence of contaminating tumour cells in the matched normal sample and/or low variant allele frequency that may result from low tumour purity.

### Somatic copy number alterations

SCNAs were identified using `ASCAT` (v3.1.2)^88^ in all autosomes and chromosome X. The required hg38 reference files for processing WES data (loci, allele, GC correction and replication timing correction files) were downloaded from https://github.com/VanLoo-lab/ascat/tree/master/ReferenceFiles/WES (git commit ID 29f2fad). Cases with a solution with a goodness-of-fit score < 0.9 were considered noisy and excluded from further analysis.

To find significant recurrent amplifications and deletions, the outputs from `ASCAT` were used to generate input files for GISTIC2 (v2.0.23)^89^, which requires segment coordinates, the number of markers and a log2-scaled copy number. For each segment obtained from segmentation from `ASCAT` analysis, the normalised depth log ratio was extracted from the `ASCAT` output files, and the number of assessed loci within each segment was used as the number of markers. Broad and focal copy number events were defined as events involving at least half a chromosome arm and less than half a chromosome arm, respectively. The residual *q*-value threshold for significant events for both broad and focal amplifications and deletions was set at 0.10. If a warning of low statistical power was given in the output, the result was not used. To remove artefactual recurrent SCNAs, we further removed `GISTIC2` focal events with an overlap of more than 40% with regions known to be problematic for sequencing and/or read mapping, as defined by the Genome in a Bottle Consortium benchmark union set of all difficult regions (v3.3; https://ftp-trace.ncbi.nlm.nih.gov/ReferenceSamples/giab/release/genome-stratifications/v3.3/GRCh38@all/Union/GRCh38_alldifficultregions.bed.gz). Finally, we excluded a focal event if less than 75% of samples predicted to have a particular SCNA in `GISTIC2` did not concur with the original `ASCAT` results.

### Prediction of chromothripsis events

We applied the definition and scoring method outlined in Vorinina *et al*.^90^ for the identification of putative chromothripsis events in tumour samples, which is based on assessing the number of CN state switches within a 50Mb region. The segmentation results from `ASCAT`, described above, were used as input. Predicted chromothripsis events were classified as high-confidence (10 or more CN state switches in 50Mb), intermediate-confidence (8 or 9 CN switches in 50Mb) or low-confidence (6 or 7 CN state switches in 50Mb). Chromothripsis events were further classified as canonical (2 or 3 different CN states) or non-canonical (> 3 CN states). For chromosome 21, which is ∼47Mb, the number of required CN state switches was scaled accordingly and rounded to the nearest whole number. For example, a scaling factor of 0.934 (46.709/50 Mb) was applied to the definitions of high-confidence (9 or more CN state switches), intermediate-confidence (7 or 8 CN state switches), and low-confidence (6 CN state switches) regions of chromothripsis.

### Identification of significantly mutated genes

To identify driver genes, we used `dNdscv` (v0.1.0; git commit ID 64f8443)^91^, which detects genes under positive selection in cancer. The reference gene annotations and covariates files used to run `dNdScv` were provided in the `dNdScv` package (RefCDS_human_GRCh38_GencodeV18_recommended.rda and covariates_hg19_hg38_epigenome_pcawg.rda, respectively). The outputs include *p*-values that were adjusted using the Benjamini-Hochberg method, and the *q*-value threshold was set at 0.10. A gene was selected if any of the *q*-values for substitutions and indels combined, substitutions only, missense mutations, nonsense mutations or indels only were below the *q*-value threshold.

### Mutational signature analysis

To identify somatic mutational signatures for single base substitutions, doublet base substitutions and indels, somatic mutations were analysed using `SigProfilerExtractor` (v1.1.21)^92^ and `SigProfilerAssignment`^93^. Results from signature decomposition and assignment were taken forward if the cosine similarity between the *de novo* extracted signatures and signatures reconstructed from assigned COSMIC signatures was >= 0.9. Similarly, on a per-sample basis, signature analysis was considered reliable if the cosine similarity between the original mutational spectrum and reconstructed spectrum was >= 0.9 and there were greater than 100 mutations per sample, per analysis type.

### Mutual exclusivity and co-occurrence of mutated genes

`DISCOVER` (r_v0.9.4)^94^ was used to test for co-occurring and mutually exclusive mutation of genes within each tumour subtype independently and within a cohort that consisted of both tumour types. A binary alteration matrix containing genes with non-silent somatic mutations was used for analysis. For both the co-occurrence and mutual exclusivity tests, the parameter selected for the estimation of the false discovery rate (FDR) was “DBH”, which implements a discrete Benjamini-Hochberg procedure that is described in Canisius *et al*.^94^. Genes used for the analysis were mutated in 2 or more samples within the cohort. A FDR threshold of 0.1 was applied.

### Analysis of whole exome germline variants

Following the Genome Analysis Toolkit (GATK) Best Practices workflow^95^, PCR duplicates from aligned WES reads in normal sample BAMs were marked using the `Picard Tools` (v2.25.4; https://broadinstitute.github.io/picard/) `MarkDuplicates` function and short variant discovery was performed using `GATK` (v4.2.6.1)^96^. Additional details are provided in the Supplementary Methods. For both SNVs and indels, only variants within 100 bp of a bait set region were retained. Functional consequences were annotated using `VEP`, as described above for the annotation of somatic variants. To identify variants present within known germline cancer predisposition genes, we looked for variants present within the set of genes used by England’s National Health Service (NHS) Cancer National Genomic Test directory (v7.2, June-2023; https://www.england.nhs.uk/wp-content/uploads/2018/08/Cancer-national-genomic-test-directory-version-7.2-June-2023.xlsx) to diagnose cancer predisposition in patients. Only variants with a moderate or high impact on the encoded protein, as predicted by `VEP`, were reported.

### Fusion gene analysis

Paired-end exome-capture transcriptome sequencing data was used to identify fusion transcripts in tumour samples. Fusion identification was performed using `STAR-Fusion` (v1.10.1)^97^, with `STAR` (v2.78a) aligner and the Trinity Cancer Transcriptome Analysis Toolkit (CTAT) genome library `StarFv1.10` for GRCh38 using GENCODE v37 (Ensembl v103) gene annotations. We used `FusionInspector` (v2.6.0) included with the STAR-Fusion package to assess the coding effect of the fusions, annotate their presence in the CTAT library and validate them *in silico* using the `STAR-Fusion` parameters `--FusionInspector validate --examine_coding_effect --denovo_reconstructFusion`. For the purpose of aiding case review of the salivary gland tumours collected, in the “discovery set” of samples we minimally filtered the fusion gene predictions, removing only fusions with less than 5 junction-spanning reads, in order to identity known fusions or putative fusions involving either *HMGA2*, *MYB*, *NFIB*, *PLAG1* or *WIF1*. To identify high-confidence fusions, we searched for fusions in the high-quality cohort (described above). A set of high-confidence fusion predictions obtained by removing those with junction reads < 5; fusions without a coding effect prediction (frameshift or inframe fusion transcirpt); fusions previously reported in normal tissues in the Genotype-Tissue Expression (GTEx) project database; and fusions involving genes described as neighbours. Lists of recurrent fusions in salivary gland tumours and frequently rearranged genes were obtain from Freiberger *et al*.^98^ and Bubola *et al*.^99^. The collation of results, filtering, summaries per cohort and plotting was performed using custom scripts.

### Pathogen identification

We used `Kraken2` (v2.1.2)^100^ to search for evidence of pathogens in the tumour samples. Briefly, unfiltered paired-end exome-capture RNA-seq FASTQ files were provided as input to `Kraken2`, which was run with a 16GB-capped reference database containing RefSeq sequences from archaea, bacteria, viruses, plasmids, humans, UniVec_Core, protozoa, fungi, and plants (https://benlangmead.github.io/aws-indexes/k2; March 2023). A confidence score threshold of 0.1 was used. Next, we calculated the proportion of minimisers from the `Kraken2` reference database found in each sample for each taxon, defined as the number of distinct minimisers out of the total clade level minimisers. To assess the significance of the results, we used a normal distribution to calculate the probability of finding *m* minimisers, out of the total *M* minimisers available for a taxon, in *n* reads of a sample, considering the total *N* reads evaluated by `Kraken2` in that sample. We used the Benjamini-Hochberg method to correct for multiple comparisons. Results with adjusted *p-*value < 0.05 were considered statistically significant.

### Shotgun cloning and Sanger sequencing

Confirmation of the *FBXW11* mutation was performed for 5 tumour samples (PD52386a, PD52405a, PD52410a, PD56543a, PD52393a) and their respective matched normal sample, where DNA stock was available (PD52386b, PD52405b, PD52393b). Validation was not performed for BCA case PD52381a due to the exhaustion of DNA stocks. The region surrounding the *FBXW11* p.F517S (g.5:171868777A>G) variant was amplified using the primers `FBXW11_F CAAAGGGAAGGTGAATTCAATCA` and `FBXW11_R TGCCCAGTTTCTCATTGTGA` in AccuStart II GelTrack PCR SuperMix with 35 cycles using DNA extracted from FFPE cores as described above. After PCR clean-up, fragments were cloned using the TOPO™ TA Cloning™ Kit and shotgun cloned into Stellar™ Competent Cells (TaKaRa). Individual colonies were picked into 96 well plate culture blocks, prepped for plasmid DNA extraction, and capillary sequenced with M13F and M13R primers. Reads were analysed by manual inspection and by alignment to the human reference genome and to exon 13 of *FBXW11*.

### Structural analysis

The crystal structure of FBXW11 (Protein Data Bank ID: 6WNX) was used to investigate the functional effect of the p.F517S substitution (Uniprot isoform E5RGC1). Wild-type FBXW11 and a model of the p.F517S protein mutant (including the water molecules present in the crystal structure) were subjected to cycles of energy-minimization, short molecular dynamics (MD) simulations (1 ns), and re-minimization with `HyperChem` (v8.0; Hypercube, Inc., Gainesville, FL). Computations were performed considering all residues within a selection sphere (20 Å from the C-alpha atom of residue 517). The Amber94 force field was used employing switched cutoffs (inner and outer radius were respectively 10 and 14 Å) and constant temperature (309.15 K). One hundred MD minimization cycles were made to produce as many conformers for both wild type FBXW11 and the F517S mutant.

### *In vitro* functional analyses

A complete list of reagents is provided in the Supplementary Methods. Abbreviations are as follows: Dulbecco’s modified Eagle’s medium (DMEM); Fetal bovine serum (FBS); cycloheximide (CHX).

Constructs. The entire coding sequence of *FBXW11* was cloned into the pcDNA6.2/V5-HisA eukaryotic expression vector (Invitrogen). The construct expressing the mutant FBXW11^F517S^ protein was generated by PCR-based site directed mutagenesis using the QuiKChange II Site-Directed Mutagenesis Kit (Agilent), as previously described^101^. All constructs were bidirectionally Sanger-sequenced for their entire open reading frame (ABI BigDye terminator Sequencing Kit v3.1) (Applied Biosystems, Foster City, CA), using a SeqStudio Genetic Analyzer (Applied Biosystems).

Cell cultures, transfections and inhibitor treatment. COS-1 and HEK293T cell lines (American Type Culture Collection, Manassas, VA) were cultured in DMEM medium supplemented with 10% heat-inactivated FBS, and antibiotics (37 °C, humidified atmosphere containing 5% CO_2_). Subconfluent COS-1 and HEK293T cells were transfected using the PEI transfection reagent according to the manufacturer’s instructions. COS-1 cells were treated with CHX (10 µg/ml) to assess protein stability. Serum-free DMEM was utilized to starve cells.

Cell lysates and co-immunoprecipitation assays. After treatment with CHX, COS-1 cells were lysed in radio-immune precipitation assay (RIPA) buffer, pH 8.0, complemented with protease and phosphatase inhibitor cocktails. Lysates were kept on ice for 30 min and then centrifuged at 16,000 x g for 30 min at 4 °C. For FBXW11/β-catenin co-immunoprecipitation (co-IP) assays, HEK293T cells were serum-starved and lysed in IP buffer containing 25 mM Tris-HCl (pH 7.4), 1% Triton X-100, 2 mM EDTA (pH 8.0), 150 mM NaCl, supplemented with protease and phosphatase inhibitors. Samples were centrifuged at 10,000 x g (20 min, 4 °C). Supernatants were collected, and their protein concentration was determined by Bradford assay, using bovine serum albumin (BSA) as a standard. In co-IP experiments, equal amounts of total proteins were immunoprecipitated using an anti-β-catenin or anti-V5 antibody cross-linked to Protein G Sepharose beads (2 h, 4 °C). The beads were recovered by centrifugation and washed six times with IP buffer. Finally, the immunoprecipitated proteins were eluted with sample buffer by incubating at 95 °C (10 min) and stored at -20 °C until immunoblotting analysis.

Immunoblotting assays were performed as previously reported^101^. In brief, cell lysates were resolved by sodium dodecyl sulfate (SDS)-polyacrylamide gel electrophoresis. Proteins were transferred to a nitrocellulose membrane using the Trans-Blot Turbo transfer system. Blots were blocked with 5% non-fat milk powder in PBS containing 0.1% Tween-20 for 1 h and incubated with specific antibodies overnight. Primary and secondary antibodies were diluted in blocking solution. Immunoreactive proteins were detected by an enhanced chemiluminescence (ECL) detection kit, according to the manufacturer’s instructions. Densitometric analysis of protein bands was performed using NineAlliance UVITEC software (UVITEC, Cambridge, UK).

### Immunohistochemistry for β-catenin

β-catenin immunohistochemistry was performed on a Leica BOND III™ stainer (Leica Biosystems) using the validated ‘IHC Protocol F’ for human tissue, according to the manufacturer’s instructions. The primary antibody was a mouse monoclonal anti-human beta-catenin antibody (Clone beta-catenin 1, M3539, Dako, Agilent) used at 1:100 dilution.

## Data availability

Sequencing data are available from the European Genome-Phenome Archive (EGA) under study accessions EGAS00001007745 and EGAS00001007746.

## Author contributions

K.W., D.J.A. and M.T. wrote the paper with assistance from all of the authors. K.W., M.D.C.W.H., J.B., S.C., V.O., I.V. and A.D. performed analysis of the next generation sequencing data. J.A.B., I.W., I.F., M.H., M.A., NdSA, C.M., A.S. and T.B. performed histopathological and immunohistochemical analysis. L.v.d.W., E.A. and K.S. managed the sample collections and the generation of the sequence data and were responsible for ensuring ethical compliance along with the pathologists who supplied the cases. M.M., E.B., G.R., A.C., M.T. performed the FBXW11 modelling analyses and functional studies.

## Ethics statement

All patients gave written informed consented to this research which was IRB approved at local ethical review boards and also by the Sanger Institute.

## Acknowledgements

This study was supported by a Medical Research Council (MRC) program grant to D.J.A. (MR/V000292/1) and an AIRC grant to M.T. (IG 28768). This research was funded in whole, or in part, by Wellcome Trust Grant 206194. The authors thank the CASM IT and Pathogens and Microbes teams, Wellcome Sanger Institute, for their assistance in the development of analysis pipelines. We also wish to thank the patients and their families.

## SUPPLEMENTARY FIGURE LEGENDS

**Supplementary Figure 1**: Review of salivary gland tumours and selected mutations and fusions. Shown are salivary gland tumours collected for this study that passed quality assurance. Tumours with and without a matched normal sample are included for case review, however, BCAC and BCA cases without a matched normal sample were excluded from the final analysis cohort. Review of cases were based on both histopathology and select somatic mutations and fusion. Fusions shown are those previously identified in salivary gland tumours (SGT fusions) and other fusions involving *HMGA2*, *WIF1*, *NFIB*, *MYB* or *PLAG1*. Exact breakpoints were identified for *MYB*::*NFIB*, *NFIB*::*MYB*, *MYB*::*AHI1* and *HMGA2*::*WIF1* fusions. Abbreviations: ACC, adenoid cystic carcinoma; EMC, epithelial myoepithelial carcinoma; ME, myoepithelioma; BCA, basal cell adenoma; BCAC, basal cell adenocarcinoma; BCAC/EMC, differential diagnosis of BCAC and EMC; CXPA, carcinoma ex-pleomorphic adenoma; PA/PA_HMGA2-WIF1, pleomorphic adenoma or pleomorphic adenoma with *HMGA2::WIF1*; PA_HMGA2, pleomorphic adenoma with a *HMGA2* fusion other than *HMGA2*::*WIF1*; PA/ME/CXPA, differential diagnosis of PA, ME and CXPA; NOS, salivary gland tumour, not otherwise specified. I35T^ indicates a mutation found by manual inspection of sequencing read alignments. Cases with a consensus BCA or BCAC diagnosis and a matched normal sample were taken forward for further analysis.

**Supplementary Figure 2:** Hematoxylin and eosin staining of a salivary gland basal cell adenoma. Basal cell adenoma (PD52375a) with a tubular and trabecular growth pattern of bilayered ducts and minimal intervening cellular stroma. The cells are bland and show no mitotic figures. The tumour border was encapsulated and smooth with no infiltration (not shown).

**Supplementary Figure 3:** Hematoxylin and eosin staining of a salivary gland basal cell adenocarcinoma. The tumour (PD56536c) shows a solid circumscribed nodule on the left and an infiltrative basaloid small nested component on the right which is infiltrating soft tissue.

**Supplementary Figure 4:** Recurrently mutated genes in salivary gland basal cell adenoma (BCA) and basal cell adenocarcinoma (BCAC). Shown are genes with protein-altering mutations in at least 2 samples in (a) BCA and (b) BCAC. TMB is tumour mutation burden, mutations/Mb. Fusions genes in red text indicate fusions found in the Trinity Cancer Transcriptome Analysis Toolkit human fusion library. Samples marked as “Discovery set only” did not pass strict criteria for transcriptome sequencing quality control (see Methods) and may have a higher false discovery rate as a result. The copy number panel indicates the tumours that had copy number loss of chromosome arms 5q and/or 16q, which was significant in the BCAC cohort.

**Supplementary Figure 5:** Decomposition of mutation signatures extracted from a cohort of salivary gland basal cell adenocarcinomas (BCACs). *De novo* signature extraction identified 1 single base substitution (SBS) signature (SBS96A) and 1 doublet base substitution (DBS) signature (ID83A) in a cohort of 9 BCAC and 2 BCAC with a differential diagnosis of epithelial-myoepithelial carcinoma (BCAC/EMC). Signature decomposition with COSMIC mutational signatures was performed, identifying SBS1, SBS5 and SBS44 in SBS96A and ID2 and ID7 in ID83A. Reconstructed SBS96A and ID83A signatures had cosine similarities of 0.946 and 0.907 with the original extracted SBS96A and ID83A signatures, respectively. In sample PD56546c, signatures found in tumours with deficient mismatch repair (SBS44, ID2 and ID7) were active. The signature decomposition and assignment statistics for PD56546c are shown.

**Supplementary Figure 6:** Somatic copy number alterations in salivary gland basal cell adenoma (BCA) and salivary gland basal cell adenocarcinoma (BCAC). (a) A chromothripsis-like event was identified on chromosome 3 of PD56545a, a tumour with differential diagnosis of BCAC and epithelial-myoepithelial carcinoma. (b) Examples of copy number alteration in BCAC in samples (b) PD56541a, a lung metastasis, and (c) PD56536c, a primary tumour from the parotid gland. (d) Copy number alterations in a BCA, PD52405a, from the parotid gland. The top panels show the allele-specific copy number, the middle panels show the log2 depth ratios and the lower panels show the b-allele frequencies (BAF) in the tumour. Plots were generated by ASCAT (see Methods).

**Supplementary Figure 7:** Location of mutations in select genes in salivary gland basal cell adenocarcinoma. Shown are lollipop plot representations of proteins and protein domains. (a) KMT2D mutations in 2 salivary gland basal cell adenocarcinoma (BCAC; red and blue), 1 BCAC with differential diagnosis of epithelial-myoepithelial carcinoma (green) and 1 basal cell adenoma (BCA). Mutations from the same tumour are indicated by colour. PHD is the plant homeodomain finger. (b) Missense mutations in 2 BCACs were located in the ubiquitin carboxyl-terminal hydrolase (UHC) domain of CYLD. (c) DICER1 mutations in 2 BCACs. Mutations from the same tumour are indicated by colour. Purple rectangles represent ribonuclease IIIa (left) and ribonuclease IIIb domains (right). (d) Two BCACs had mutations affecting the same amino acid, p.K171M and p.K171T. Pkinase is the protein kinase domain. Plots were generated using MutationMapper on the cBioPortal website (https://www.cbioportal.org/mutation_mapper).

**Supplementary Figure 8:** Germline variants in the salivary gland basal cell adenoma (BCA) and basal cell adenocarcinoma (BCAC). Shown are protein-altering germline variants in (a) BCA (b) BCAC. Genes shown are those included in the National Health Service England’s National Genomic Test Directory for somatic and inherited cancers (v7.2) that were either not found in the gnomAD database (v3.1) or present with an overall population frequency < 0.0001. Variants annotated in the ClinVar database (20230121) as benign or likely benign are not shown.

