## Supplementary Materials for "Activation of Wnt/β-catenin signalling by mutually exclusive *FBXW11* and *CTNNB1* hotspot mutations drives salivary gland basal cell adenoma"

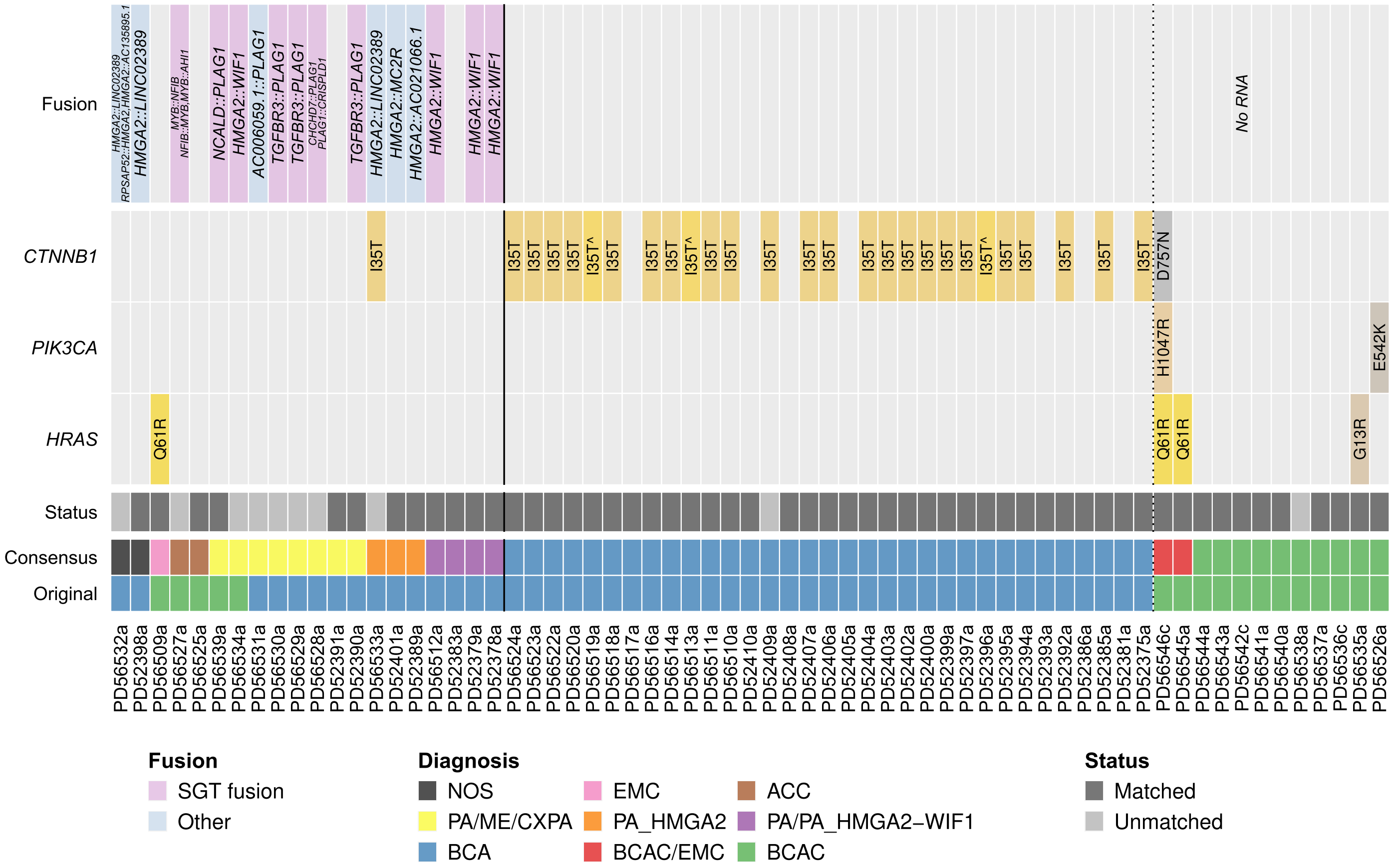


**Supplementary Figure 1**: Review of salivary gland tumours and selected mutations and fusions. Shown are salivary gland tumours collected for this study that passed quality assurance. Tumours with and without a matched normal sample are included for case review, however, BCAC and BCA cases without a matched normal sample were excluded from the final analysis cohort. Review of cases were based on both histopathology and select somatic mutations and fusion. Fusions shown are those previously identified in salivary gland tumours (SGT fusions) and other fusions involving *HMGA2*, *WIF1*, *NFIB*, *MYB* or *PLAG1*. Exact breakpoints were identified for *MYB*::*NFIB*, *NFIB*::*MYB*, *MYB*::*AHI1* and *HMGA2*::*WIF1* fusions. Abbreviations: ACC, adenoid cystic carcinoma; EMC, epithelial myoepithelial carcinoma; ME, myoepithelioma; BCA, basal cell adenoma; BCAC, basal cell adenocarcinoma; BCAC/EMC, differential diagnosis of BCAC and EMC; CXPA, carcinoma ex-pleomorphic adenoma; PA/PA_HMGA2-WIF1, pleomorphic adenoma or pleomorphic adenoma with HMGA2::WIF1; PA_HMGA2, pleomorphic adenoma with a HMGA2 fusion other than HMGA2::WIF1; PA/ME/CXPA, differential diagnosis of PA, ME and CXPA; NOS, salivary gland tumour, not otherwise specified. I35T^ indicates a mutation found by manual inspection of sequencing read alignments. Cases with a consensus BCA or BCAC diagnosis and a matched normal sample were taken forward for further analysis.

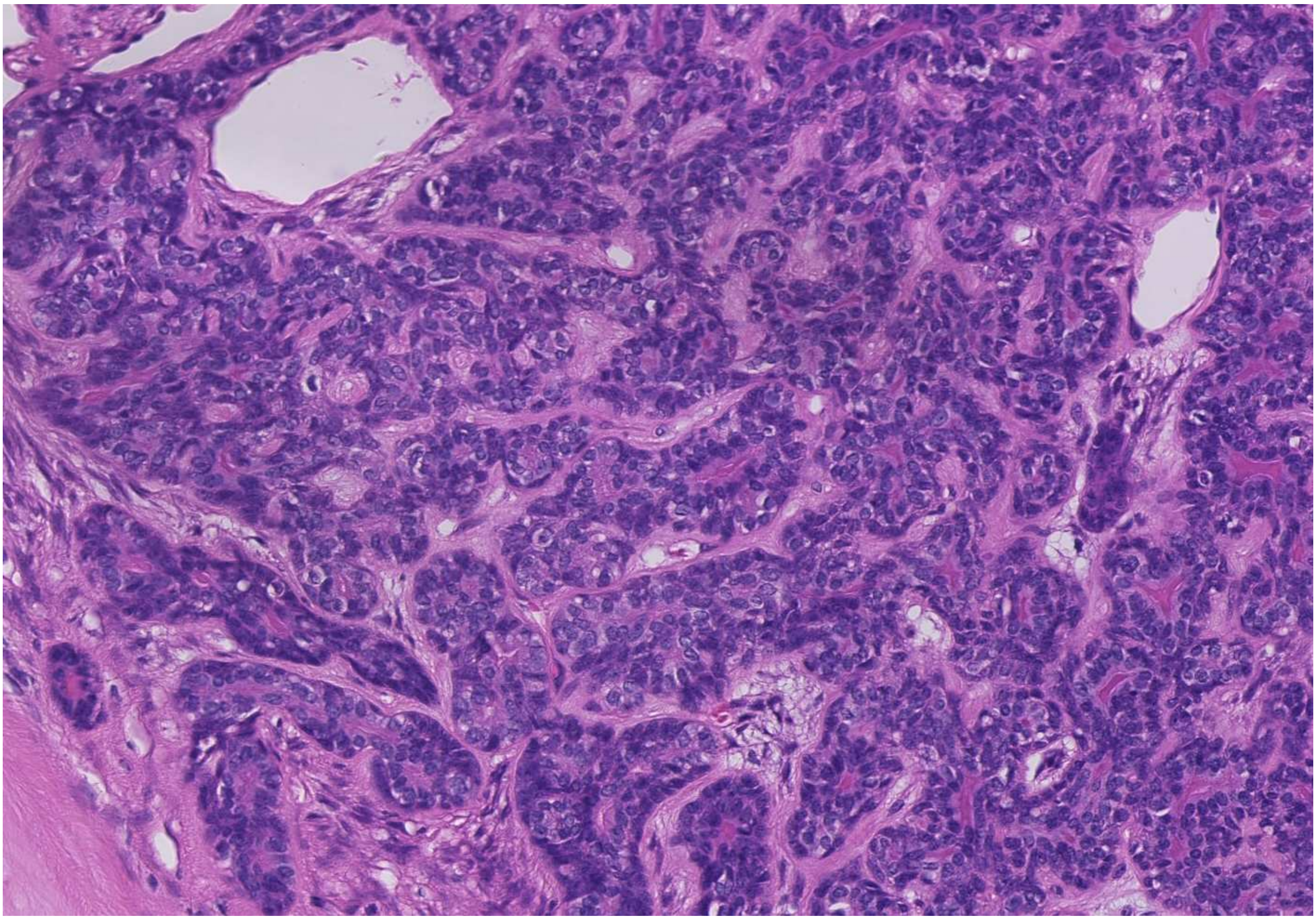

**Supplementary Figure 2:** Hematoxylin and eosin staining of a salivary gland basal cell adenoma. Basal cell adenoma (PD52375a) with a tubular and trabecular growth pattern of bilayered ducts and minimal intervening cellular stroma. The cells are bland and show no mitotic figures. The tumour border was encapsulated and smooth with no infiltration (not shown).

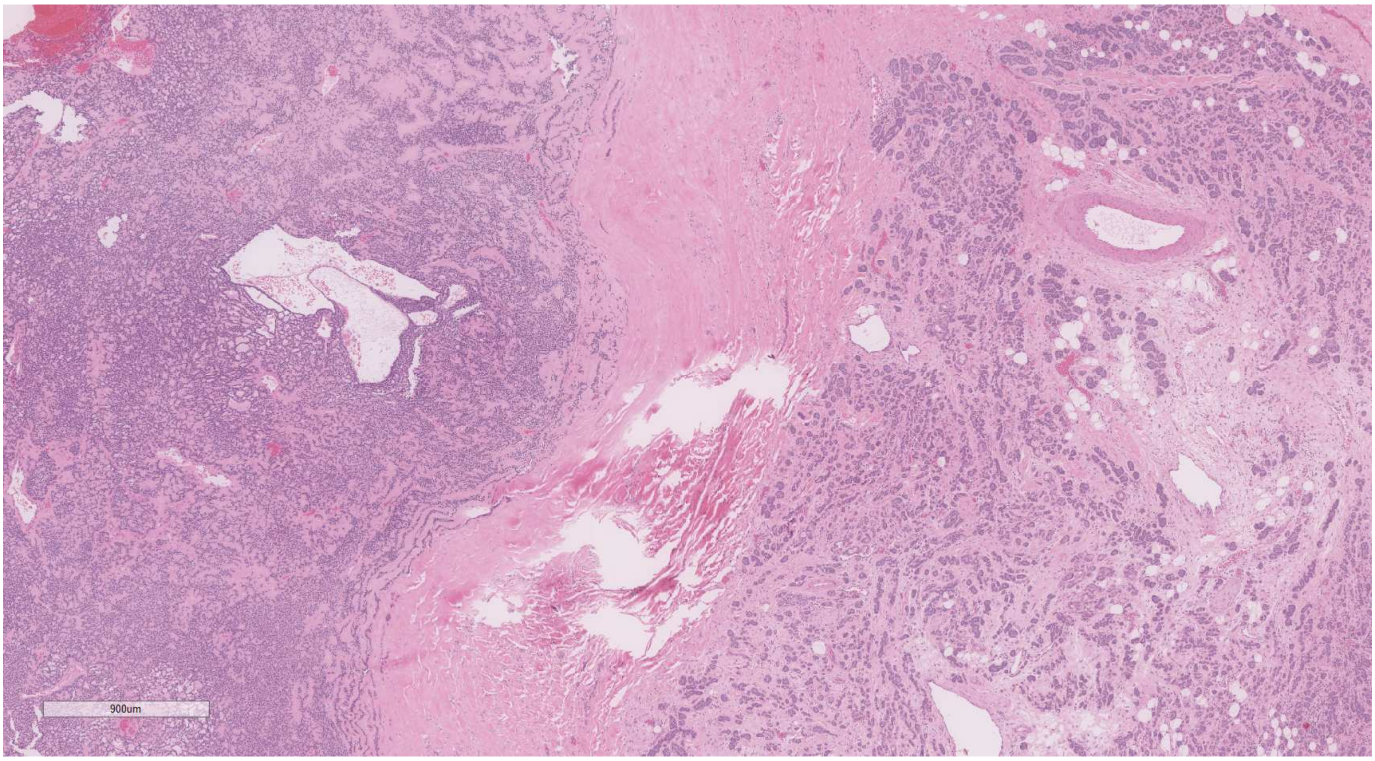


**Supplementary Figure 3:** Hematoxylin and eosin staining of a salivary gland basal cell adenocarcinoma. The tumour (PD56536c) shows a solid circumscribed nodule on the left and an infiltrative basaloid small nested component on the right which is infiltrating soft tissue.

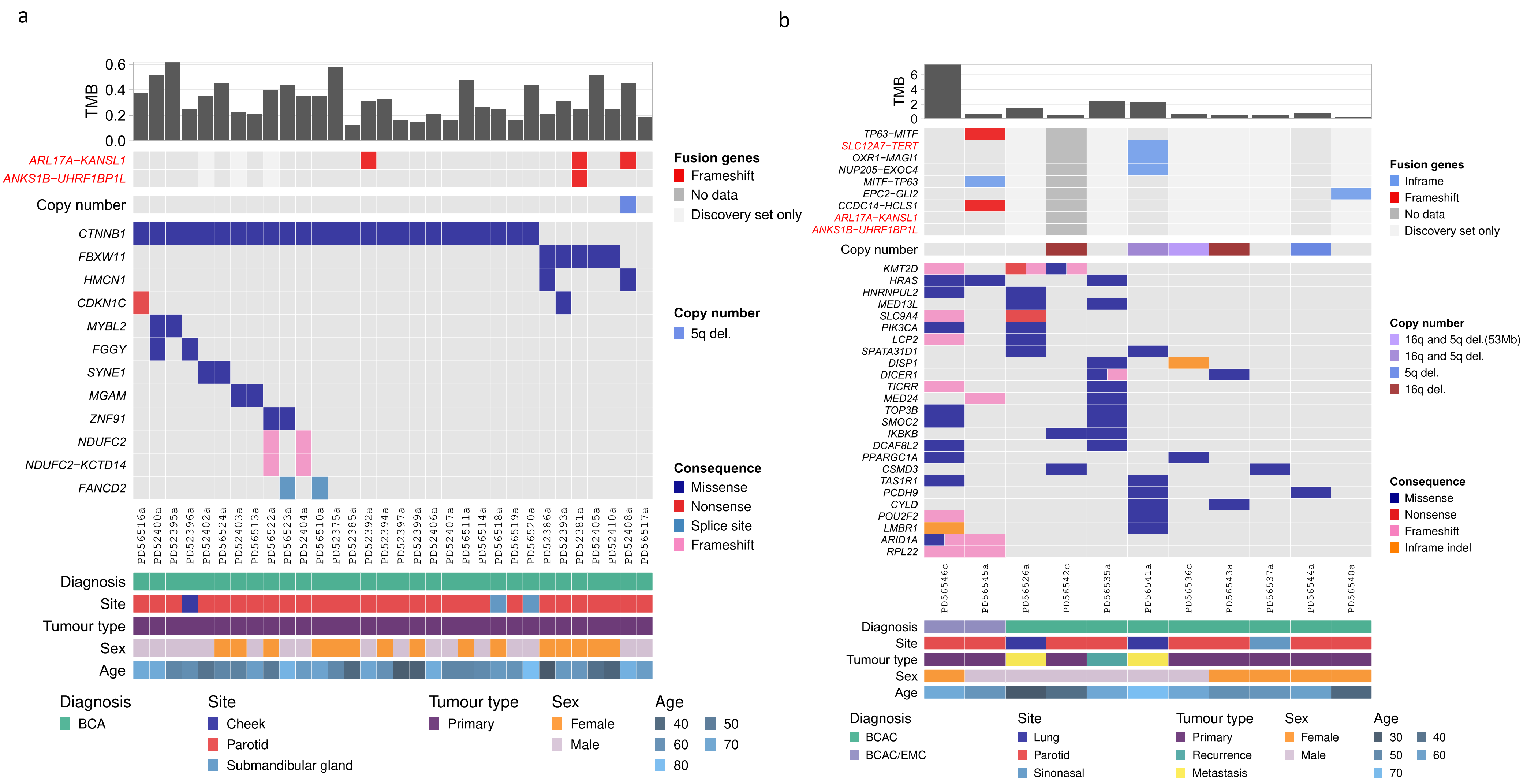


**Supplementary Figure 4:** Recurrently mutated genes in salivary gland basal cell adenoma (BCA) and basal cell adenocarcinoma (BCAC). Shown are genes with protein-altering mutations in at least 2 samples in (a) BCA and (b) BCAC. TMB is tumour mutation burden, mutations/Mb. Fusions genes in red text indicate fusions found in the Trinity Cancer Transcriptome Analysis Toolkit human fusion library. Samples marked as "Discovery set only" did not pass strict criteria for transcriptome sequencing quality control (see Methods) and may have a higher false discovery rate as a result. The copy number panel indicates the tumours that had copy number loss of chromosome arms 5q and/or 16q, which was significant in the BCAC cohort.

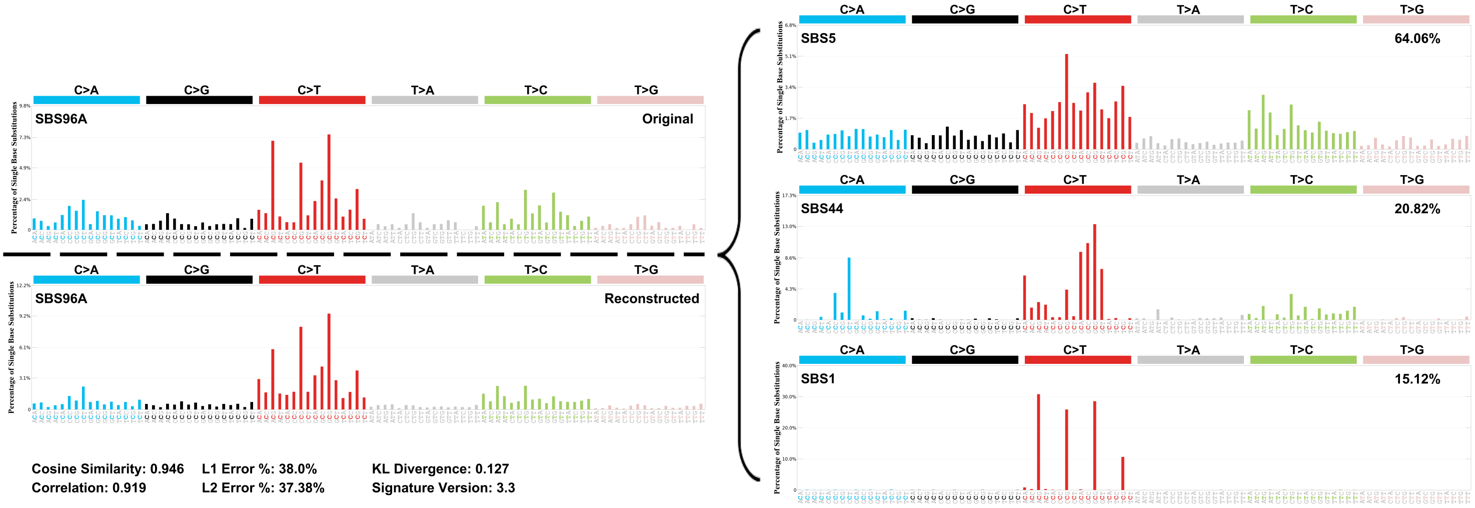


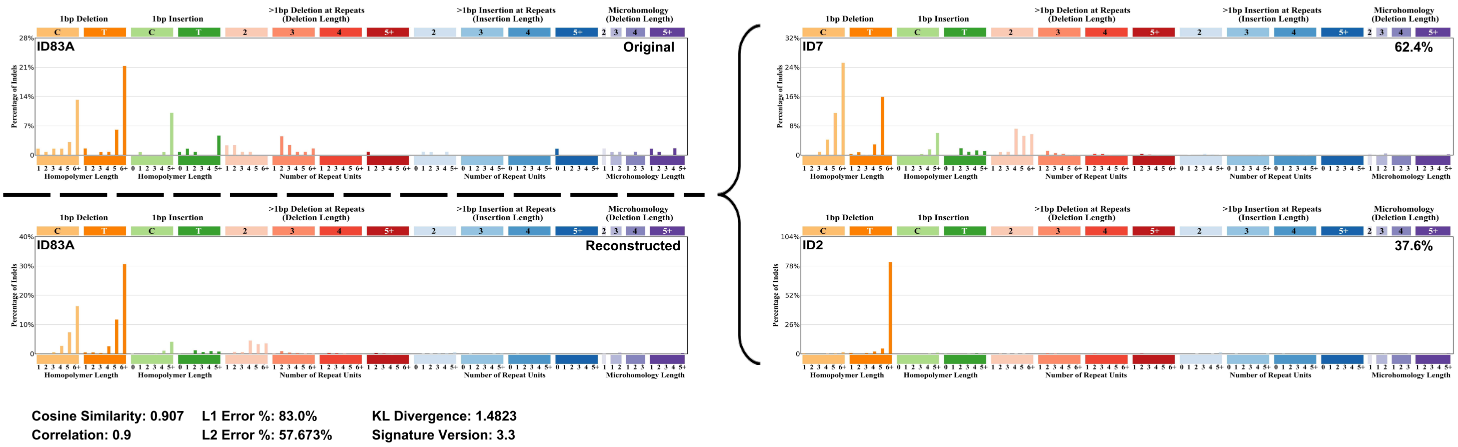


| Signature activities in PD56546c | | | |
|---|---|---|---|
| SBS1 | SBS44 | ID2 | ID7 |
| 62 | 200 | 34 | 54 |

| Solution statistics for PD56546c | | | | | | | | |
|---|---|---|---|---|---|---|---|---|
| Signature type | Mutations | Cosine similarity | L1 norm | L1 Norm % | L2 norm | L2 norm % | KL divergence | Correlation |
| SBS | 263 | 0.869 | 164.4 | 62.7 | 35.1 | 62.9 | 0.59 | 0.846 |
| ID | 88 | 0.944 | 49.9 | 56.7 | 11.4 | 33.7 | 0.52 | 0.939 |

**Supplementary Figure 5:** Decomposition of mutation signatures extracted from a cohort of salivary gland basal cell adenocarcinomas (BCACs). *De novo* signature extraction identified 1 single base substitution (SBS) signature (SBS96A) and 1 doublet base substitution (DBS) signature (ID83A) in a cohort of 9 BCAC and 2 BCAC with a differential diagnosis of epithelial-myoepithelial carcinoma (BCAC/EMC). Signature decomposition with COSMIC mutational signatures was performed, identifying SBS1, SBS5 and SBS44 in SBS96A and ID2 and ID7 in ID83A. Reconstructed SBS96A and ID83A signatures had cosine similarities of 0.946 and 0.907 with the original extracted SBS96A and ID83A signatures, respectively. In sample PD56546c, signatures found in tumours with deficient mismatch repair (SBS44, ID2 and ID7) were active. The signature decomposition and assignment statistics for PD56546c are shown.

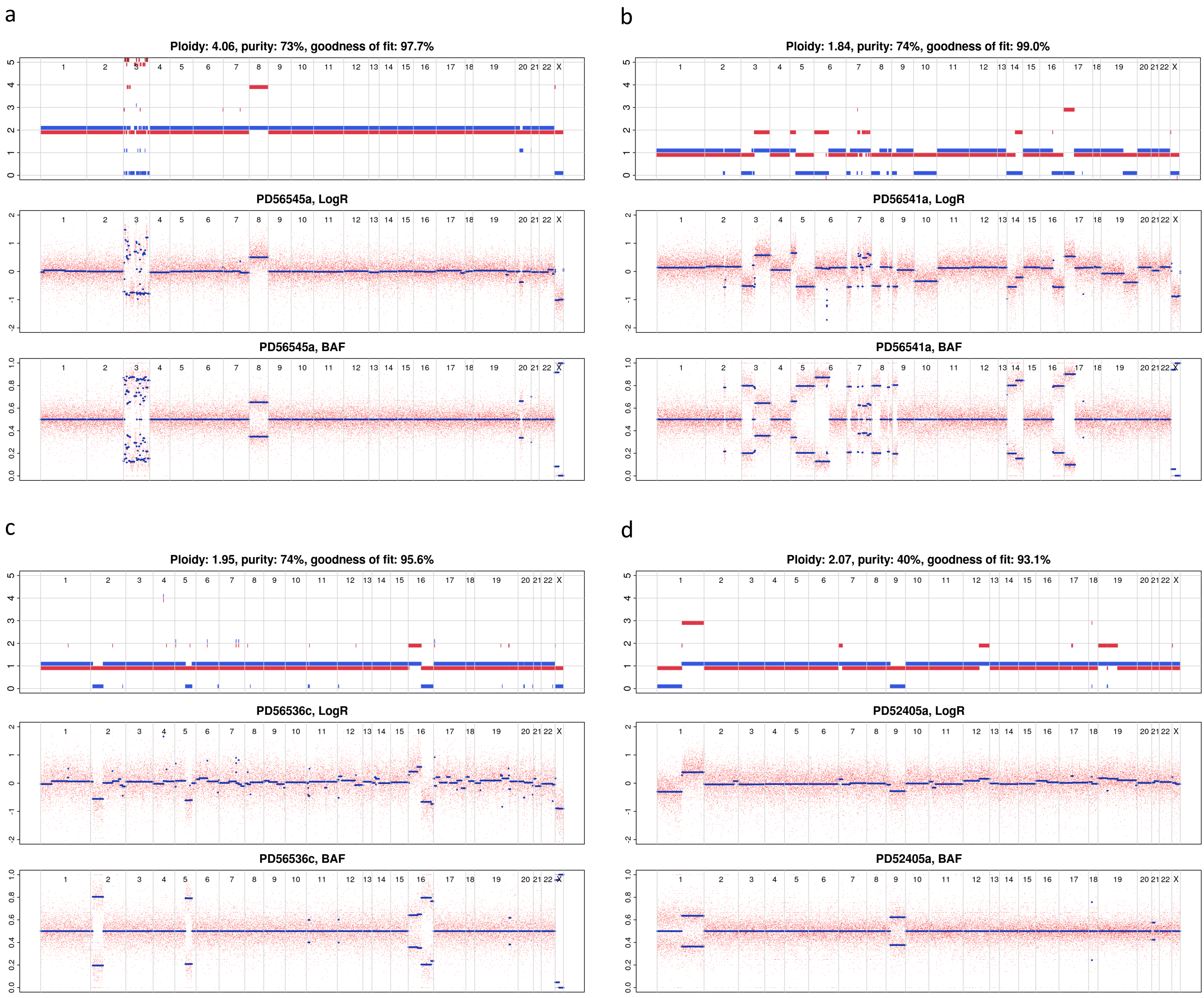


**Supplementary Figure 6:** Somatic copy number alterations in salivary gland basal cell adenoma (BCA) and salivary gland basal cell adenocarcinoma (BCAC). (a) A chromothripsis-like event was identified on chromosome 3 of PD56545a, a tumour with differential diagnosis of BCAC and epithelial-myoepithelial carcinoma. (b) Examples of copy number alteration in BCAC in samples (b) PD56541a, a lung metastasis, and (c) PD56536c, a primary tumour from the parotid gland. (d) Copy number alterations in a BCA, PD52405a, from the parotid gland. The top panels show the allele-specific copy number, the middle panels show the log2 depth ratios and the lower panels show the b-allele frequencies (BAF) in the tumour. Plots were generated by ASCAT (see Methods).

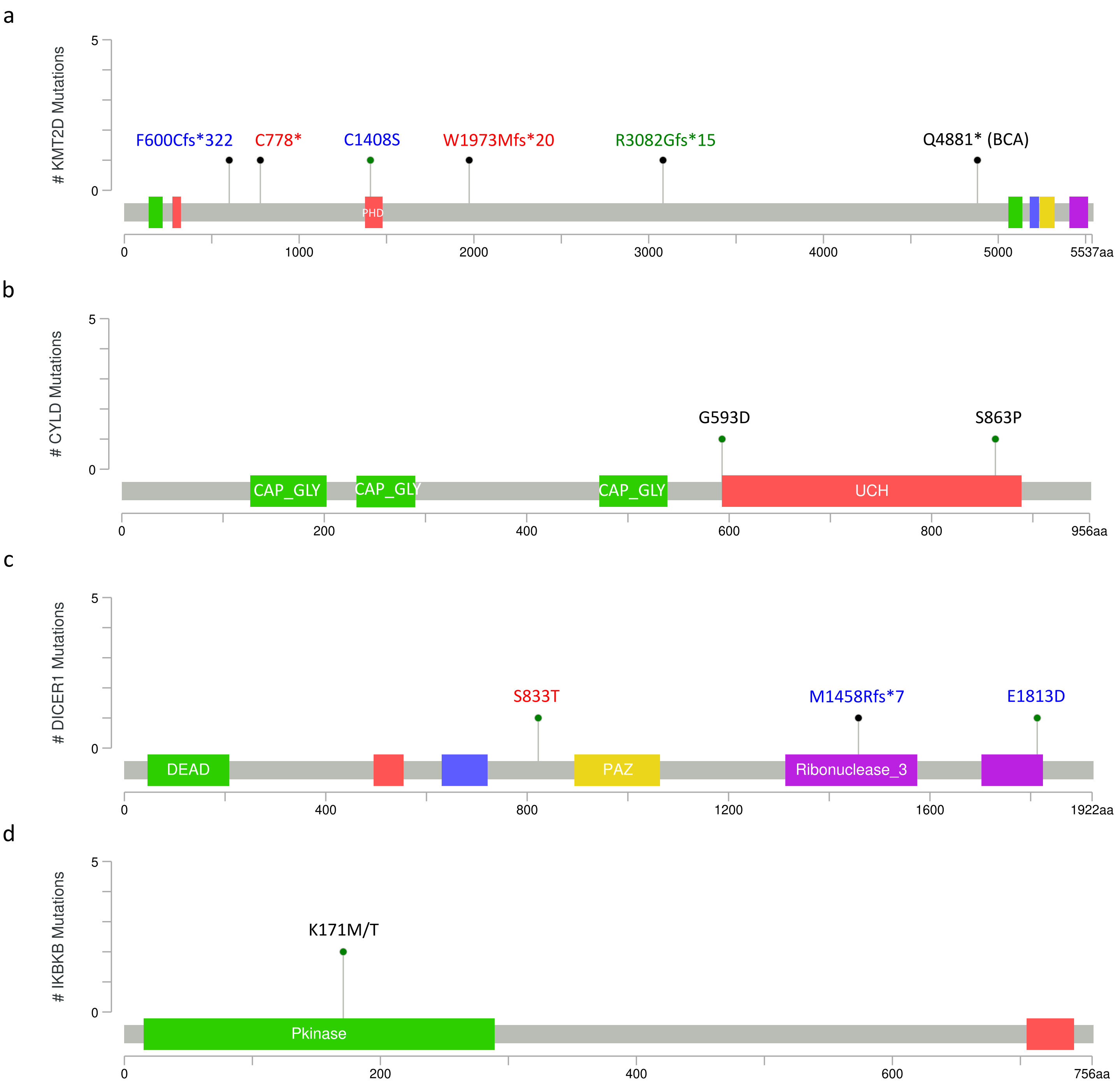


**Supplementary Figure 7:** Location of mutations in select genes in salivary gland basal cell adenocarcinoma. Shown are lollipop plot representations of proteins and protein domains. (a) KMT2D mutations in 2 salivary gland basal cell adenocarcinoma (BCAC; red and blue), 1 BCAC with differential diagnosis of epithelial-myoepithelial carcinoma (green) and 1 basal cell adenoma (BCA). Mutations from the same tumour are indicated by colour. PHD is the plant homeodomain finger. (b) Missense mutations in 2 BCACs were located in the ubiquitin carboxyl-terminal hydrolase (UHC) domain of CYLD. (c) DICER1 mutations in 2 BCACs. Mutations from the same tumour are indicated by colour. Purple rectangles represent ribonuclease IIIa (left) and ribonuclease IIIb domains (right). (d) Two BCACs had mutations affecting the same amino acid, p.K171M and p.K171T. Pkinase is the protein kinase domain. Plots were generated using MutationMapper on the cBioPortal website (https://www.cbioportal.org/mutation_mapper).

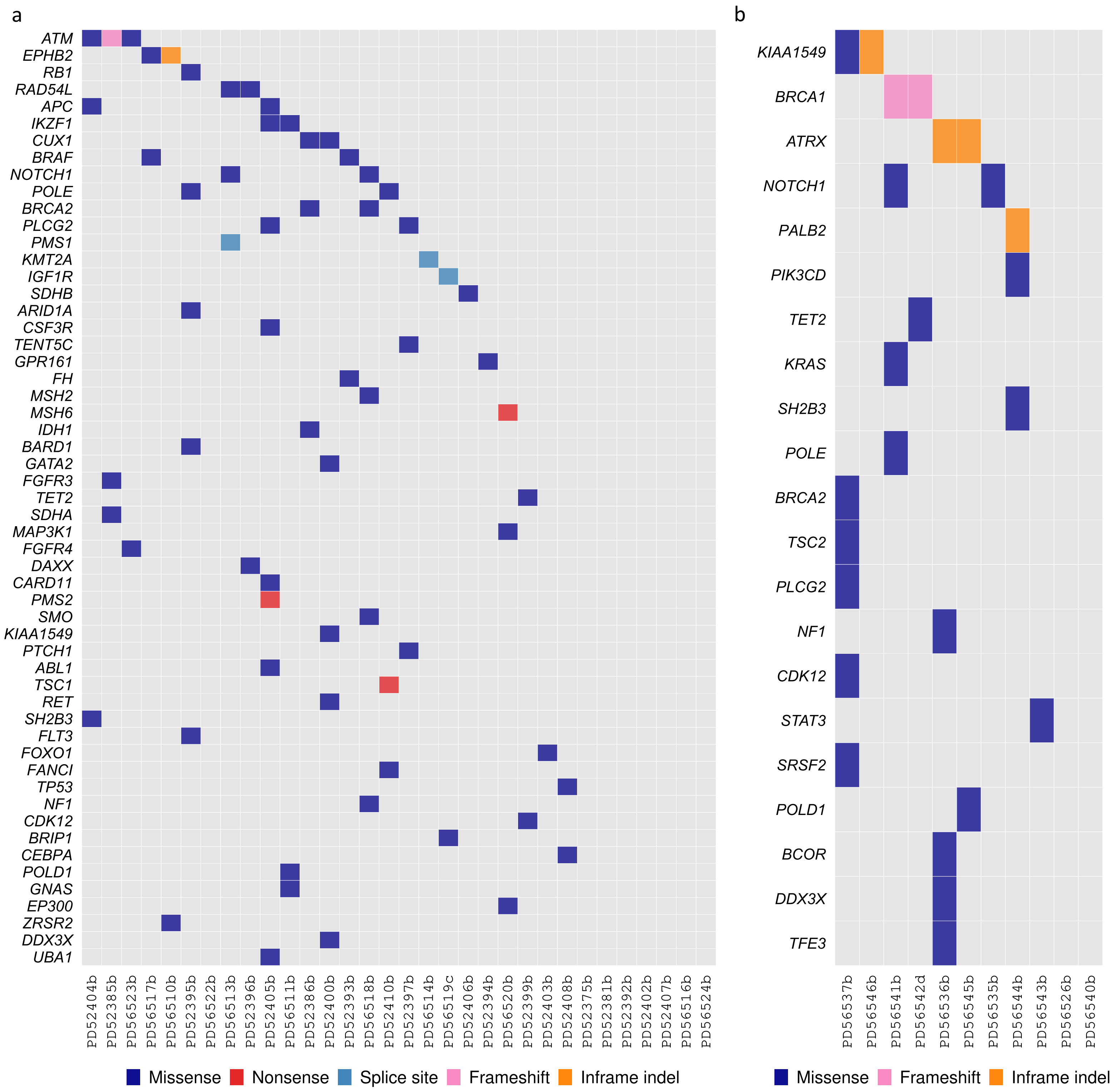


**Supplementary Figure 8:** Germline variants in the salivary gland basal cell adenoma (BCA) and basal cell adenocarcinoma (BCAC). Shown are protein-altering germline variants in (a) BCA (b) BCAC. Genes shown are those included in the National Health Service England's National Genomic Test Directory for somatic and inherited cancers (v7.2) that were either not found in the gnomAD database (v3.1) or present with an overall population frequency < 0.0001. Variants annotated in the ClinVar database (20230121) as benign or likely benign are not shown.

**Supplementary Table 1: Sample metadata.** Shown are samples collected and sequenced for this study. Samples from the same patient have the same numerical prefix. “DNA QC status” indicates whether the DNA samples passed quality control checks for depth, contamination and tumour-normal sample concordance, if applicable. “RNA QC status” indicates whether the transcriptome sequencing data passed quality control checks. If the transcriptome sequencing data was available but quality control requirements were not met, the data was used for fusion discovery only. The “Diagnosis” column is the consensus diagnosis after review and “Original diagnosis” is the diagnosis provided at the time of sample collection. The tumour types are: P, primary; R, recurrence; M, metastasis. Sex is: F, female; M, male. The “Included in cohort” column indicates inclusion or exclusion from the final BCA and BCAC cohorts, which comprised of paired tumour-normal pairs only. Transcriptome data was excluded if the corresponding DNA data was also excluded.

**See Excel spreadsheet Supp_Table_1_SG_BCA_BCAC_metadata.xslx**

**Supplementary Table 2: Somatic variants identified from whole-exome sequencing of salivary gland tumours with matched normal tissue.** Somatic variants in (a) salivary gland basal cell adenoma, (b) salivary gland basal cell adenocarcinoma and (c) cases excluded after review of histopathology and genetic profiling. Variants are provided in mutation annotation format (MAF; https://docs.gdc.cancer.gov/Data/File_Formats/MAF_Format/), with additional information from variant annotation with the Ensembl Variant Effect Predictor (VEP, v103), dbSNP (v155), ClinVar (release 20230121) and the gnomAD (v3.1) database (see Methods).

**See Excel spreadsheet: Supp_Table_2_SG_BCA_BCAC_variants.xslx**

**Supplementary Table 3: Salivary gland basal cell adenoma (BCA) and basal cell adenocarcinoma (BCAC) tumour mutation burden.** Mutation rates (mutations/Mb) were calculated from the BCA and BCAC cohorts, primary BCAC only, recurrent (rec.) or metastatic (met.) cases only and all BCAC, excluding BCAC with differential diagnosis of EMC (BCAC/EMC).

| Mutation rate (mutations/Mb) | BCA | BCAC (all) | BCAC (primary) | BCAC (rec./met.) | BCAC (excluding BCAC/EMC) |
|---|---|---|---|---|---|
| Samples | 32 | 11 | 8 | 3 | 9 |
| Mean | 0.32 | 1.59 | 1.42 | 2.05 | 1.04 |
| Median | 0.31 | 0.68 | 0.61 | 2.30 | 0.66 |
| Range | 0.12-0.62 | 0.22-7.48 | 0.22-7.48 | 1.45-2.38 | 0.22-2.38 |

**Supplementary Table 4: Gene fusions identified in salivary gland basal cell adenoma (BCA) and basel cell adenocarcinoma (BCAC).** Fusions were identified from analysis of transcriptome sequencing data with `STAR-fusion`, `fusionInspector` and `Trinity`, and annotated with `fusionAnnotator` (see Methods). Shown are predicted fusions that have at least 5 junction spanning reads and annotated as either inframe of frameshift fusions. Fusions in red are present in one or more fusion databases in the Trinity Cancer Transcriptome Analysis Toolkit genome library StarFv1.10.

**See Excel spreadsheet: Supp_Table_4_Gene_Fusions.xlsx**

**Supplementary Table 5: Significantly mutated genes in salivary gland basal cell adenoma and basal cell adenocarcinoma.** The `dndscv` algorithm (see Methods) was used to identify significantly mutated genes in salivary gland basal cell adenoma (BCA) and basal cell adenocarcinoma (BCAC) cohorts. Shown are genes with *q*-values < 0.1 (in bold) when considering substitutions only, indels only, and substitutions and indels together (global *q*-value). All *q*-values were obtained using the Benjimini-Hochberg multiple testing correction.

| Gene | Cohort | Substitution q-value | Indel q-value | Global q-value |
|---|---|---|---|---|
| *CTNNB1* | BCA | **0** | 1.0 | 0 |
| *FBXW11* | BCA | **1.7e-06** | 1 | **4.0e-05** |
| *KMT2D* | BCAC | 1.0 | **0.028** | **0.012** |
| *HRAS* | BCAC | **0.012** | 1.0 | **0.09** |
| *RPL22* | BCAC | 1.0 | **0.028** | 0.25 |

**Supplementary Table 6: Sanger sequencing validation of a recurrent *FBW11* mutation.** PCR amplification, shotgun cloning, and Sanger sequencing of both tumour and matched normal samples, where available, was performed to validate a recurrent mutation in exon 13 of *FBXW11* on chromosome 5 at position 171868777 (A>G) (c.1550T>C or p.F517S on transcript ENST00000517395.6 and protein ENSP00000428753.2, respectively). The mutation was present in 14-60% of tumour clones sequenced (*n* = 10-15 clones per sample) in 4 BCAs and 1 BCAC with the mutation and absent in the 3 matched normal samples for which we had DNA stocks (*n* = 6-14 clones per sample), which indicated that the mutation was indeed present and somatic. Additionally, the p.I526M mutation in sample PD52393a was also validated (Figure 3). Validation was not performed for the 6th BCA case, PD52381a, due to a lack of DNA stocks; however, the mutation was present in the transcriptome of this sample.

| Sample | Tissue type | Clones with mutation (%) |
|---|---|---|
| PD52386a | Tumour | 6/14 (42.9%) |
| PD52386b | Normal | 0/8 (0.0%) |
| PD52393a | Tumour | 6/10 (60.0%) |
| PD52393b | Normal | 0/14 (0.0%) |
| PD52405a | Tumour | 2/14 (14.2%) |
| PD52405b | Normal | 0/6 (0.0%) |
| PD52410a | Tumour | 5/15 (33.3%) |
| PD56543a | Tumour | 4/15 (26.7%) |

**Supplementary Table 7: Genes in the Wnt signaling pathway.** Somatic variant lists from the salivary gland basal cell adenoma and basal cell adenocarcinoma cohorts were queried for mutations in the genes listed below, which are part of the Wnt/β-catenin signalling pathway.

| **Gene** | **Hugo Symbol (Ensembl v103)** |
|---|---|
| *Axin1* | *AXIN1* |
| *Axin2* | *AXIN2* |
| *APC* | *APC* |
| *CDK14* | *CDK14* |
| *RNF43* | *RNF43* |
| *ZNRF3* | *ZNRF3* |
| *YAP* | *YAP1* |
| *TAZ* | *WWTR1* |
| *GSK3-β* | *GSK3B* |
| *CK1-α* | *CSNK1A1* |
| *CTNNB1* | *CTNNB1* |
| *β-TrCP* | *BTRC* |
| *LRP5* | *LRP5* |
| *LRP6* | *LRP6* |
| *Dvl1* | *DVL1* |
| *Dvl2* | *DVL2* |
| *Dvl3* | *DVL3* |
| *CBP* | *CRBBP* |
| *p300* | *EP300* |
| *BRG1* | *SMARCA4* |
| *BCL9* | *BCL9* |
| *Pygo1* | *PYGO1* |
| *Pygo2* | *PYGO2* |
| *FBXW7* | *FBXW7* |
| *β-TrCP2* | *FBXW11* |

**Supplementary Table 8: Significant recurrent copy number alterations.** Shown are significant focal and broad somatic copy number alterations (SCNAs) from `GISTIC2` analysis (see Methods), with residual *q*-value < 0.1 (Benjamini-Hochberg method). Significant focal SCNAs were compared to the Genome in a Bottle (GIAB) consortium's difficult regions benchmarking set and removed from analysis if the fraction of overlap with difficult regions was >= 0.4. Finally, for each sample with an amplification or deletion in a significant SCNA, `ASCAT` copy number calls were evaluated for concordance (see Methods). Any `GISTIC2` SNCA was rejected if there was less than 75% concordance with `ASCAT` calls. The number of samples included in the `GISTIC2` analysis was 32 and 11 for the BCA and BCAC cohorts, respectively. CGC, COSMIC Cancer Gene Census; TSG, tumour suppressor gene; N/A, not applicable.

| **Cohort** | **Chromosome region** | **Amplification or deletion** | **Residual *q*-value** | **Comment** | **Fraction overlap with difficult regions** | **Concordance with ASCAT, % (Samples in agreement)** |
|---|---|---|---|---|---|---|
| BCA | chr14:22088606-22506783 (0.42 Mb) | Amp | 0.022 | *TRA*/*TRD* locus | 0.10 | 100 (4/4) |
| BCA | chr5:140675698-141526846 (0.85 Mb) | Del | 0.0048 | Protocadherin-β gene cluster | 0.21 | 100 (2/2) |
| BCAC | chr2:11154272-29192809 (18.0 Mb) | Del | 0.084 | CGC TSGs *DNMT3A* and *ASXL2* are in this region | 0.12 | 100 (3/3) |
| BCAC | chr 5q | Del | 0.097 | No mutations in CGC genes in samples with 5q deletion. *APC* and *FBXW11* are on 5q | N/A | 100 (2/2) |
| BCAC | chr 16q | Del | 8.6xe-07 | *CYLD* (a CGC TSG) mutations found in 2 cases with 16q deletion (PD56541a, PD56543a) | N/A | 100 (4/4) |

**Supplementary Table 9: Selected germline variants in patients with salivary gland basal cell adenoma (BCA) and basal cell adenocarcinoma (BCAC).** Variants were identified in mismatch repair genes (*MSH6*, *PMS2* and *MSH2*) and *BRCA* genes. The genomic position is relative to the reference genome GRCh38. The rsID is the dbSNP (v155) rsID; gnomAD AF is the total population allele frequency of the variant in the gnomAD (v3.1) database; ClinVar interpretation is the ClinVar database (release 20230121) clinical significance of the variant on disease; Impact is the Variant Effect Predictor (v103) predicted effect of the variant on the protein; COSMIC variant indicates whether the variant has been reported in the COSMIC database; Sample is the BCA or BCAC sample with the variant; Somatic mutation rate is the tumour mutation burden given in mutations/Mb. Variants in the genes below with a ClinVar clinical significance of 'benign' or 'likely benign' were excluded.

| Gene | Genomic position | Amino acid change | rsID | gnomAD AF | ClinVar interpretation | Impact | COSMIC variant | Sample | Somatic mutation rate (mut./Mb) |
|---|---|---|---|---|---|---|---|---|---|
| *MSH6* | 2:47798646 | p.E221D | rs41557217 | 6.3e-04 | Conflicting interpretations | Moderate | COSV52286342 (2 samples) | PD52394b (BCA) | 0.33 |
| *MSH6* | 2:47806641 (C>T) | p.R1331* | rs267608094 | 6.6e-06 | **Pathogenic** | High | COSV52278149 (3 samples) | PD56520b (BCA) | 0.44 |
| *PMS2* | 7:5986883 (G>A) | p.R628* | rs63750451 | N/A | **Pathogenic** | High | N/A | PD52405b (BCA) | 0.52 |
| *MSH2* | 2:47463105 (C>G) | p.D487E | rs35107951 | 9.8e-05 | Conflicting interpretations | Moderate | N/A | PD56518b (BCA) | 0.25 |
| *BRCA1* | 17:43057062 (T>TG INS) | p.Q1777Pfs*74 | rs80357906 | 5.3e-05 | **Pathogenic** | High | N/A | PD56541b (BCAC) | 2.3 |
| *BRCA1* | 17:43092848 (GTT>G DEL) | p.K894Tfs*8 | rs8035797 | 6.6e-06 | **Pathogenic** | High | COSV99066399 (2 samples) | PD56542d (BCAC) | 0.46 |
| *BRCA2* | 13:32337456 (T>C) | p.I1034T | rs545974734 | 1.3e-05 | Uncertain significance | Moderate | N/A | PD52386b (BCA) | 0.21 |
| *BRCA2* | 13: 32363253 (A>G) | p.K2684R | rs80359043 | N/A | Uncertain significance | Moderate | N/A | PD52386b (BCA) | 0.21 |
| *BRCA2* | 13:32376724 (G>A) | p.R2896H | rs80359128 | 6.6e-06 | Conflicting interpretations of pathogenicity | Moderate | COSV66452365 (2 samples) | PD56518b (BCA) | 0.25 |
| *TSC1* | 19:132903668 (C>A) | p.E731* | rs397514820 | N/A | **Pathogenic** | High | N/A | PD52410b (BCA) | 0.25 |
| *EP300* | 22:41169525 (G>A) | p.D1399N | rs1057519889 | N/A | **Likely pathogenic** | Moderate | COSV54326888 (54 samples) | PD56520b (BCA) | 0.44 |

### Supplementary Methods

The following sections provide additional details to those provided in the Methods in the main text.

#### Additional methods for transcriptome sequencing quality control

To select samples for a high-quality analysis cohort, we discarded any samples that reported: expression profiling efficiency < 40%; 3' bias < 0.3 or 3' bias > 0.5; proportion of reads intersecting rRNAs > 2.5%; < $20 \times 10^7$ read pairs; a higher sum total of low quality, ambiguous and multi-aligned reads than total reads counted; or number of genes with five counts or more < $14 \times 10^3$.

#### Additional methods for cgpCaVEMan variant calling and flagging

Below are `cgpCaVEMan` parameters for tumours with matched normal samples:

```
  -reference
$REF_BASE/GRCh38_full_analysis_set_plus_decoy_hla.fa.fai \
  -outdir $OUTDIR \
  -tumour-bam $TUM_BAM \
  -normal-bam $NORM_BAM \
  -ignore-file $REF_BASE/genome.gap.tab \
  -tum-cn-default 5 \
  -norm-cn-default 2 \
  -species Human \
  -species-assembly GRCh38 \
  -flag-bed-files $REF_BASE/caveman/flagging \
  -germline-indel $GERM_INDEL \
  -unmatched-vcf $REF_BASE/caveman/unmatched_vcf_dir \
  -seqType exome \
  -tumour-protocol WSX \
  -normal-protocol WSX \
  -normal-contamination 0.1 \
  -noflag \
  -flagConfig $REF_BASE/caveman/flag.vcf.config.ini \
  -flagToVcfConfig $REF_BASE/caveman/flag.to.vcf.convert.ini \
  -threads $CPU_TO_USE
```

CaVEMan flagging was run after casmsmartphase was used to identify MNVs, as described in the Methods in the main text. The following parameters were used for flagging:

```
cgpFlagCaVEMan.pl \
      --input $VCF_IN \
      --outFile $VCF_OUT \
      --species Human \
      --reference \
      --studyType WXS \
```

```
      --tumBam $TUM_BAM \
      --normBam $NORM_BAM \
      --bedFileLoc $REF_BASE/caveman/flagging \
      --unmatchedVCFLoc $REF_BASE/caveman/unmatched_vcf_dir \
      --annoBedLoc $REF_BASE/vagrent/e103 \
      --flagConfig $REF_BASE/caveman/flag.vcf.config.ini
   --flagToVcfConfig $REF_BASE/caveman/flag.to.vcf.convert.ini
```

The `--annoBedLoc` option had to be provided for the script to run, however, we used Ensembl VEP (v103; see main Methods and below) to predict variant consequences.

For tumours without a matched normal sample, an '*in silico*' BAM file was used in place of a normal BAM file and –normal-contamination was set to 0.0, as described in the Methods section of the manuscript.

Additional parameters, flags and BED files applied to the SNV and MNV calling and flagging in the `flag.vcf.config.ini` file were as follows:

```
[HUMAN_WXS PARAMS]
keepSW=0
minAnalysedQual=11
maxMatchedNormalAlleleProportion=0.03
maxPhasingMinorityStrandReadProportion=0.04
readPosBeginningOfReadIgnoreProportion=0.08
readPosTwoThirdsOfReadExtendProportion=0.08
pentamerMinPassAvgQual=20
samePosMaxPercent=80
maxTumIndelProportion=10
maxNormIndelProportion=10
minPassAvgMapQual=21
minPassPhaseQual=30
minDepthQual=30
minNormMutAllelequal=30
minRdPosDepth=8
vcfUnmatchedMinMutAlleleCvg=3
vcfUnmatchedMinSamplePct=5
matchedNormalMaxMutProportion=0.20
minSingleEndCoverage=10
depthCutoffProportion=0.5
maxCavemanMatchedNormalProportion=0.2
withinXBpOfDeletion=10
minGapPresentInReads=20
minMeanMapQualGapFlag=10
minGapFlagDistEndOfReadPercent=75
maxGapFlagDistFromEndOfReadProp=0.13

[HUMAN_WXS FLAGLIST]
flagList=<<LST
```

```
depthFlag
readPositionFlag
matchedNormalFlag
pentamericMotifFlag
avgMapQualFlag
centromericRepeatFlag
codingFlag
snpFlag
phasingFlag
tumIndelDepthFlag
sameReadPosFlag
hiSeqDepthFlag
annotationFlag
unmatchedNormalVcfFlag
singleEndFlag
matchedNormalProportion
alignmentScoreReadLengthAdjustedFlag
clippingMedianFlag
alnScoreMedianFlag
cavemanMatchNormalProportionFlag
withinGapRangeFlag
LST

[HUMAN_WXS BEDFILES]
centromericRepeatBed=centromeric_repeats.bed.gz
simpleRepeatBed=simple_repeats.bed.gz
snpBed=snps.bed.gz
annotatableBed=gene_regions.bed.gz
codingBed=codingexon_regions.sub.bed.gz
germlineIndelBed=germline_indel.bed
highSeqDepthBed=genome.tab.gz
```

The `centromeric_repeats.bed` and `simple_repeats.bed` files were generated using the UCSC Table Browser, as described here:

https://www.ncbi.nlm.nih.gov/pmc/articles/PMC6097606/#S16
https://www.ncbi.nlm.nih.gov/pmc/articles/PMC6097605/#S14

except GRCh38 was used instead of GRCh37.

The `codingBed` and `annotatableBed` files were not used, as we were not using `VAGrENT` to annotate the variant calls.

The `snps.bed.gz` file contains a manually curated list of SNPs from dbSNP. The SNPs were manually curated to remove oncogenic variants.

The `genome.tab.gz` file contains regions with extreme sequencing depth (more than 8 standard deviations from the mean depth in an internal reference set of BAMs), which are excluded from variant calling. Genic regions are excluded from the `genome.tab.gz` file.

Germline indels were not used for filtering, however, a non-empty BED file was required. Therefore, a tab-delimited BED file with "`1 0 1`" was used as input.

An unmatched normal panel consisting of sequencing data from normal tissue of 98 individuals was used to filter both polymorphisms and artefacts that occur from sequencing and read alignment artefefacts, using the `--unmatchedNormalVcfFlag` option. Generation of this panel is described here:

https://www.ncbi.nlm.nih.gov/pmc/articles/PMC6097605/).

A detailed description of CaVEMan flags is available here:

https://github.com/cancerit/cgpCaVEManPostProcessing/wiki/flags-and-settings

To identify MNVs, adjacent SNVs were first extracted from VCFs generated by cgpCaVEMan using the `casmsmartphase` (v0.1.8) '`generate-bed`' utility (https://github.com/cancerit/CASM-Smart-Phase/releases/tag/0.1.8) with the `--markhz` option to generate a BED file for SmartPhase. SmartPhase was run with the parameters: `-m 0 -x -g adjacent_snvs.bed`. The `casmsmartphase` '`merge-mnvs`' utility was then used to select MNVs by selecting adjacent MNVs with the *cis* phasing and a minimal confidence score cutoff of 0.1 (`--exclude 30 --cutoff 0.1`) and produce a modified cgpCaVEMan VCF with MNVs merged into a single VCF entry. Adjacent homozygous SNVs identified by '`generate-bed`' were also merged into MNVs. Variants were flagged using the `cgpcavemanpostprocessing` (v1.10) `cgpFlagCaVEMan.pl` utility using 'WXS' mode for exomes.

**Additional methods for `cgpPindel` variant calling and flagging**

The following parameters were used to run `cgpPindel`:

```
  -o $WORK/$pair \
  -r $REF_BASE/GRCh38_full_analysis_set_plus_decoy_hla.fa \
  -t $TUM_BAM \
  -n $NORM_BAM \
  -s $REF_BASE/pindel/simpleRepeats.bed.gz \
  -f $REF_BASE/pindel/WSX_Rules.lst \
  -g $REF_BASE/vagrent/e${ENSM_VER}/codingexon_regions.indel.bed.gz
\
  -u $REF_BASE/pindel/pindel_np.v5.gff3.gz \
  -st WXS \
  -e chrUn%,HLA%,%_alt,%_random,chrM,chrEBV \
  -b $REF_BASE/shared/HiDepth_mrg1000_no_exon_coreChrs_v3.bed.gz \
  -sf $REF_BASE/pindel/softRulesFragment.lst \
```

```
-noflag
-c $CPU_TO_USE
```

The `HiDepth_mrg1000_no_exon_coreChrs_v3.bed.gz` file is the same as the genome.gap.tab.gz file described above, but the former has 1-based start positions, and the later has 0-based start positions.

The `pindel_np.v5.gff.gz` file consists of indels found in a panel of unmatched normal samples used to filter out polymorphisms and artefacts from sequencing and read mapping mis-alignment. The generation of this file is described here:

https://www.ncbi.nlm.nih.gov/pmc/articles/PMC6097606/#S16

Although VAGrENT annotation was not used, a `codingexon_regions.indel.bed.gz` file must be provided for flagging. The file was created from Ensembl v103 genes models, as described here:

https://currentprotocols.onlinelibrary.wiley.com/doi/10.1002/0471250953.bi1508s52.

The flagging rules (`WXS_Rules.lst` file) applied to the indel calls were as follows:

```
FF001
FF002
FF003
FF004
FF005
FF006
FF007
FF019
FF020
```

The definitions of these flags are available here:

https://github.com/cancerit/cgpPindel/wiki/VcfFilters

Soft flagging with FF017 was also applied (`softRulesFragment.lst)`, which indicates overlap with a simple repeat. However, variants were not hard filtered based on this flag.

To obtain library insert size statistics used as input to `cgpPindel`, `bam_stats` (v.5.6.1) was run on each sample BAM file to generate a *.bas file.

**Ensembl VEP options**

Ensembl `VEP` (v103) was run to predict the impact of germline and somatic variants. The following `VEP` options were used:

```
--db_version 103
```

```
-t SO
--format vcf
-o $output_vcf
--cache
--dir $vep_cache_path
--buffer 20000
--species homo_sapiens
--offline
--symbol
--biotype
--vcf
--sift s
--no_stats
--assembly GRCh38
--flag_pick_allele_gene
--canonical
--hgvs
--shift_hgvs 1
--fasta GRCh38_full_analysis_set_plus_decoy_hla.fa
--compress_output bgzip
--mane
--numbers
--polyphen p
--domain
--show_ref_allele
--protein
--transcript_version
$custom
```

where

```
$custom="--custom $cosmicfile,Cosmic,vcf,exact,0,CNT --custom
$clinvarfile,ClinVar,vcf,exact,0,CLNSIG,CLNREVSTAT --custom
$dbsnpfile,dbSNP,vcf,exact,0, --custom
$gnomadfile,gnomAD,vcf,exact,0,FLAG,AF"
```

and

`$gnomadfile` is a VCF with variants from gnomAD database release v3.1.2, with the `INFO` column containing only the `FLAG` and `AF` for use with `VEP` custom annotation. For example:

```
#CHROM    POS    ID    REF    ALT    QUAL    FILTER    INFO
chr1    10031    .    T    C    .    AC0;AS_VQSR
FLAG=AC0,AS_VQSR;AF=0
chr1    10037    .    T    C    .    AS_VQSR
FLAG=AS_VQSR;AF=2.60139e-05
```

The original VCFs were downloaded from:

https://gnomad.broadinstitute.org/downloads

`$clinvarfile` is a VCF from ClinVar release (dated 20230121. The file was downloaded from:

https://ftp.ncbi.nlm.nih.gov/pub/clinvar/vcf_GRCh38/weekly/clinvar_20230121.vcf.gz

`$dbsnpfile` is a VCF with variants from dbSNP release v155. For compatibility with `VEP`, chromosome names were renamed from RefSeq chromosome accessions (e.g. NC_000001.11) to chromosome numbers (e.g. chr1). The original VCF was downloaded from:

https://ftp.ncbi.nih.gov/snp/archive/b155/VCF/dbSNP155.GRCh38.GCF_000001405.39.vcf.gz

`$cosmicfile` is a VCF with COSMIC v97 coding and non-coding variants (GRCh38, normalised), with the `INFO` column containing the sample counts. For example:

```
#CHROM   POS     ID           REF   ALT   QUAL   FILTER   INFO
chr1     10108   COSV70831266  C     T     .      .        CNT=1
chr1     10151   COSV70830383  T     A     .      .        CNT=1
chr1     10175   COSV70830377  T     A     .      .        CNT=1
chr1     10181   COSV70830549  A     T     .      .        CNT=2
```

The original files were downloaded from https://cancer.sanger.ac.uk/cosmic.

**Additional details for somatic copy number analysis**

The required hg38 reference files for processing WES data using `ASCAT` (loci, allele, GC correction and replication timing correction files) were downloaded from https://github.com/VanLoo-lab/ascat/tree/master/ReferenceFiles/WES (git commit ID 29f2fad). The loci and allele files were used as input to the `ascat.perpareHTS` function, along with tumour and matched normal BAM files. Other parameters used were as follows: `genomeVersion = 'hg38', minCounts = 10, min_base_qual = 20, min_map_qual = 35` and `seed = 485028101` for reproducibility. A BED file containing the genomic coordinates of the exome pull-down regions sequenced was also provided, as recommended. The gender parameter was determined by the clinical data provided for each patient, and alleleCount (v.4.3.0; https://github.com/cancerit/alleleCount) was also used. The outputs of `ascat.prepareHTS` were used to run the `ascat.correctLogR` function with the reference GC and replication timing files followed by `ascat.aspcf`, which was then run with `penalty = 70` and `seed = 483024451` for reproducibility. Finally, the `ascat.runAscat` function was run using `gamma = 1` to estimate purity, ploidy and allele-specific copy number at each loci.

`GISTIC2` was run with the following parameters: `-refgene hg38.UCSC.add_miR.160920.refgene.mat -genegistic 1 -smallmem 1 -broad 1 -brlen 0.75 -conf 0.95 -armpeel 1 -savegene 1 -gcm extreme -v 20 -`

ta 0.25 -td 0.25. The required hg38.UCSC.add_miR.160920.refgene.mat file is included with the GISTIC2 package.

**Additional details for mutational signature analysis**

SigProfilerExtractor performs *de novo* extraction of mutational signatures and assigns known signatures to samples by refitting known COSMIC signatures to the extracted signatures using SigProfilerAssignment. SigProfilerExtracter was run in exome mode using GRCh38 as the reference and opportunity genome, 500 replicates for non-negative matrix factorisation (NMF) with 1 to 10 signatures. The solution implemented for downstream analysis was the optimal solution provided by SigProfilerExtractor (referred to as the 'suggested solution').

**Additional details for germline variant calling**

Variants were first called in each sample using the GATK HaplotypeCaller function in –ERC GVCF mode with parameters -G StandardAnnotation, -G StandardHCAnnotation, -G AS_StandardAnnotation, followed by the creation of a genomicsdb database using GenomicsDBImport. Joint genotyping was then performed using GenotypeGVCFs using default parameters. For hard-filtering of variants, SNVs and indels were separated using the Picard Tools (v2.27.1) SelectVariants function and filtered using Picard Tools (v2.27.1) VariantFiltration. For SNVs, the following parameters were used for variant filtration: QUAL<30.0, SOR>3.0, FS>60, MQ <40, MQRankSum < -12.5 and ReadPosRankSum< -8.0, and for indel filtration: QD <2.0, QUAL <30.0, FS >200.0 and ReadPosRankSum <-20.0.

**Additional details for pathogen identification using Kraken2**

Kraken2 was executed with the following options: --paired --gzip-compressed --use-names --confidence 0.1 --db path/to/DB --report path/to/report --report-minimizer-data --output /dev/null path/to/fastq1 path/to/fastq2. To generate MPA-style (MetaPhlAn) outputs, the --use-mpa-style option was used in place of --report-minimizer-data.

**Additional details for *in vitro* functional analyses**

Reagents. Dulbecco's modified Eagle's medium (DMEM), antibiotics and prestained protein SHARPMASS VI markers were obtained from Euroclone (Wetherby, UK). Fetal bovine serum (FBS), and ECL Western Blotting Detection reagents were from Thermo Fisher Scientific (Waltham, MA). Phosphate-buffered saline (PBS) was purchased from Capricorn Scientific (Ebsdorfergrund, Germany). pcDNA6.2/V5-HisA eukaryotic expression vector was obtained from Invitrogen (Carlsbad, CA). The human β-catenin pcDNA3 plasmid was a gift from Eric Fearon (Addgene plasmid #16828; http://n2t.net/addgene:16828; RRID:Addgene_16828). QuiKChange II Site-Directed Mutagenesis kit was obtained from Stratagene (La Jolla, CA). Polyethylenimine (PEI) transfection reagent was purchased from Polysciences (Warrington, PA). Protease and phosphatase inhibitor cocktails and cycloheximide (CHX) were from Sigma-

Aldrich (St. Louis, MO). Protein G Sepharose was obtained from GE Healthcare (Freiburg, Germany). Trans-Blot Turbo Transfer Packs were obtained from Bio-Rad Laboratories (Hercules, CA). The following antibodies were used: mouse monoclonal anti-V5 (Invitrogen); mouse monoclonal anti-β-catenin (Abcam, Cambridge, UK); mouse monoclonal anti-FLAG (Sigma-Aldrich), mouse polyclonal anti-polyubiquitin (Enzo Life Sciences, Farmingdale, NY); mouse monoclonal anti-GAPDH (Santa Cruz Biotechnology, Dallas TX); horseradish peroxidase conjugated anti-mouse or anti-rabbit (Thermo Fisher Scientific).
